# A *De Novo* Design Strategy to Convert FAcD from Dimer to Active Monomer

**DOI:** 10.1101/2025.01.11.632520

**Authors:** Ke-Wei Chen, Cui-zhen Wang, Hui-si Huang, Xing Wu, Jia-Nan Chen, Jia-He Qiu, Kin Kuan HOI, Tian-Yu Sun, Jian-bo Wang, Yun-Dong Wu

## Abstract

Enzymes largely exist in various oligomeric states, but monomeric enzymes are more conducive to industrial applications. Converting an oligomeric enzyme into an active monomer is a significant challenge. In this study, we present a *de novo* design strategy to convert fluoroacetate dehalogenase (FAcD) from its native dimeric form to an active monomer. Using the AI-based method ProteinMPNN, we identified critical protein-protein interaction (PPI) sites at the dimer interface. ArDCA, another AI tool, was employed to pinpoint catalytic hotspots. Six mutants, **Mu1-Mu6**, were designed. Molecular dynamics (MD) simulations, coupled with mass spectrometry, confirmed that these mutants form stable monomers. The pre-reaction state (PRS) model predicted that three of these mutants exhibited catalytic activity. In particular, **Mu5** with 11 mutations from the wild-type, was predicted to have high catalytic activity, and was subsequently confirmed by kinetics experiment, with a *k*_cat_ of 672.2 min^-1^ and a T_50_^30^ > 100 °C, comparable to the wild-type enzyme (*k*_cat_ = 676.3 min^-1^, T_50_^30^ = 84 °C). Notably, the Y149M mutation increased catalytic activity nearly forty-fold, demonstrating the effectiveness of our design strategy.

## Main

Enzyme catalysis has long been recognized as a cornerstone of biochemical processes, offering unmatched specificity, efficiency, and mild reaction conditions compared to traditional chemical catalysts^1^. In recent years, key advances such as directed evolution^2^, enzyme immobilization^3^, multi-enzymatic cascades^4^, and the integration of biocatalysis with chemical catalysis^5^ have significantly broadened the scope and efficacy of enzymatic processes, enabling them to meet the growing demands of green chemistry and industrial biotechnology. In particular, enzyme catalysis has demonstrated remarkable potential in environmental, energy, and pharmaceutical applications, such as the degradation of polyethylene terephthalate (PET) plastic^6–8^, artificial CO_2_ fixation^9^, and the synthesis of bioactive compounds like the anti-cancer drug vinblastine^10^ and the diabetes medication sitagliptin^11^. Enzyme catalysis stands as a genuinely green and sustainable technology, offering not only significant environmental benefits but also cost-effectiveness, driven by ongoing advancements in biotechnology.

Prior research has shown that most enzyme proteins are oligomeric, predominantly as dimers. Dimeric enzymes often exhibit unique cooperative effects, such as substrate binding at one active site enhancing substrate binding, catalysis, or product release at the neighboring subunit^12^. These inter-subunit cooperative effects can be achieved through various mechanisms, including pronounced conformational asymmetry^13^, inter-subunit proton wire^14–17^, electron transfer^18^, or heat capacity reduction^19, 20^. However, in some enzymes, the active form is a monomer, although higher-order forms do exist^21–23^. In some functional proteins, the monomeric form can reduce innate immune response, while dimerization can induce self-inhibition^24^, and inhibit tumor growth^25, 26^. Notably, the majority of industrially applied enzymes are small monomeric enzymes, typically ranging from 30 to 50 kDa (e.g., lipases, laccases, acylases, and proteases). Larger proteins, on the other hand, often clog pores, significantly reducing immobilization yields^27^. Furthermore, oligomeric enzymes tend to dissociate into inactive forms at elevated temperatures^28^, whereas monomeric enzymes remain stable and retain activity, thereby exhibiting superior thermostability (**Fig. 1a**)^29^. From another perspective, converting oligomeric enzymes into monomeric forms is essential for fundamental research to elucidate catalytic mechanisms and explore their catalytic potential for enhanced industrial applications^30^.

**Figure 1.**
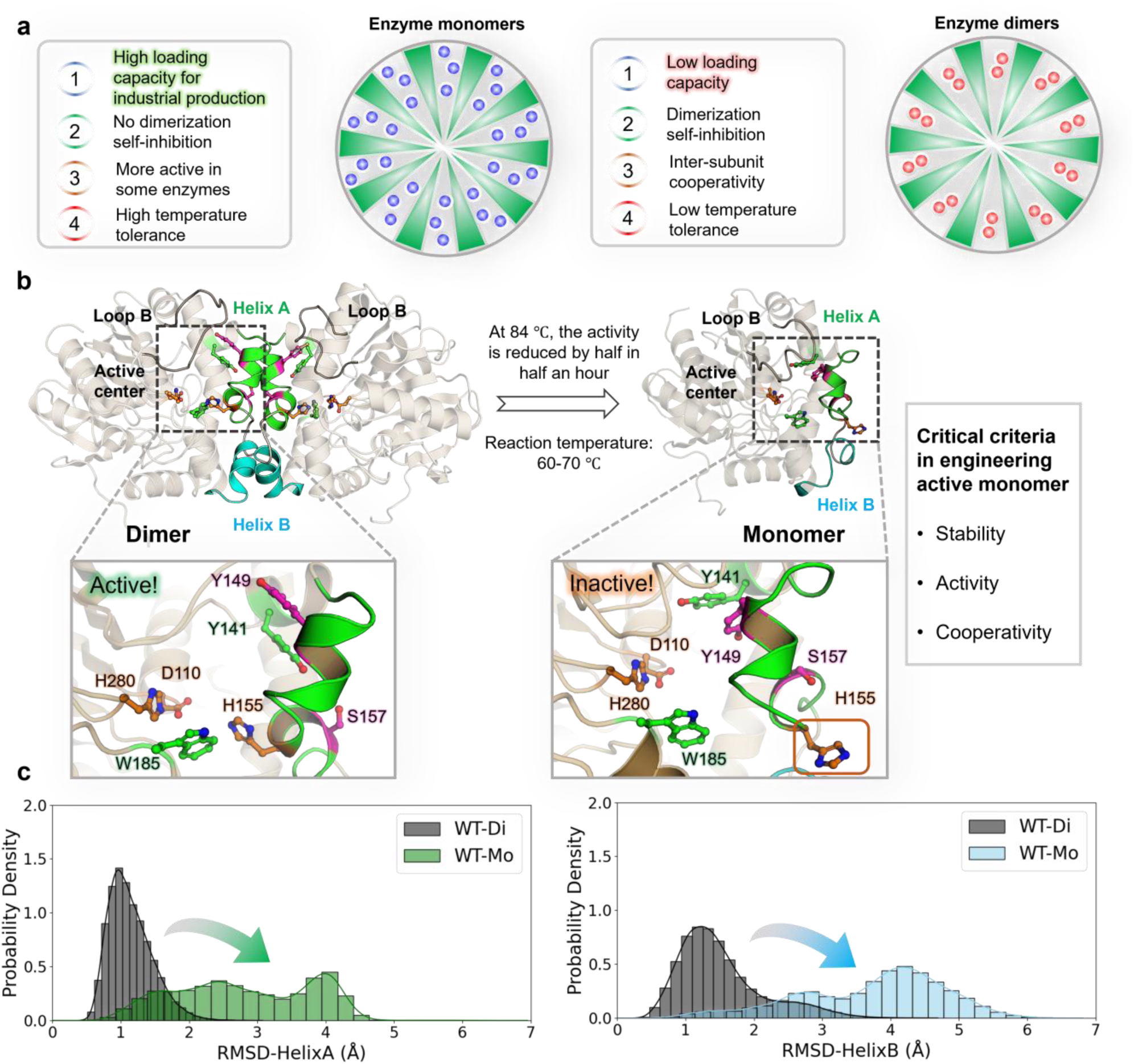
Industrial applications of enzyme monomers and characteristics of FAcD monomers. **a**, Advantages of monomers for Industrial production. **b**, Main conformations of the FAcD interface at different temperatures, with Helix A, Helix B, and catalytic triad (D110-H155-H280) labeled. **c**, Root-Mean-Square Deviation (RMSD) of the Helix A and Helix B in the active dimer and inactive monomer by molecular dynamics simulations.

Fluoroacetate dehalogenase (FAcD) is an active homodimeric enzyme that catalyzes the conversion of fluoroacetate to glycolate^31–34^. FAcD is capable of breaking the C–F bond, which holds significant promise for addressing environmental fluoride pollution, especially polyfluorinated compounds^35^. Using time-resolved crystallography, Pai et al. revealed a distinctive “half-of the sites” phenomenon in FAcD, where only one catalytic subunit (**C**) participates in catalysis during a single catalytic cycle, while the non-catalytic subunit (**NC**) can bind substrate but remains catalytically inactive^33^. Further investigations demonstrated that the substrate in the **NC** subunit plays a crucial role in modulating the conformational dynamics of gated residues W185 and Y141 in the **C** subunit, thereby enhancing catalysis and facilitating product release. This cooperativity is mediated by interfacial water chains, with key contributions from S157 and Y149 located at the dimer interface^36^. Experiments indicate that its enzyme activity diminishes at temperatures exceeding 84°C. Molecular dynamics (MD) simulations suggest that elevated temperatures lead to dissociation of the FAcD dimer into inactive monomers. The native dimer interface, including Helix A (residues M145–A160) and Helix B (residues L165–D175), is highly unstable in a monomer (**Fig. 1b, c**), rendering the monomer inactive. Converting FAcD from a dimer to a stable, active monomer could significantly enhance its thermal tolerance while facilitating improved immobilization capacity for industrial applications.

Designing oligomers to functional monomers remains exceptionally challenging. Enzyme monomer design has been extensively explored, with a primary focus on disrupting interface interactions and rational design^30, 37^. However, successful cases remain rare^38–41^. This challenge primarily stems from the need to precisely disrupt protein-protein interface interactions (PPI) without destroying active sites and cooperativity.

Here, we present a monomer design strategy that enables the conversion of FAcD dimer into an active monomer. This approach employs ProteinMPNN^42^ to identify PPI sites within the dimer interface and stabilize the enzyme monomer. Subsequently, we utilize ArDCA^43^ to identify potential active sites on the interface. By excluding these interface-active sites, we successfully converted the FAcD dimer into a functional monomer. The mutated monomer retained enzymatic activity comparable to the wild-type dimer while exhibiting enhanced thermal stability. Our strategy represents a novel attempt to design oligomeric enzymes into functional monomers, opening new avenues for protein engineering and industrial biocatalysis.

## Results

### *De novo* design FAcD monomer with ProteinMPNN and ArDCA

Rationally converting protein high-order aggregates into active monomers requires precise identification of PPI sites without destroying their activity. Although we can directly identify some key PPI sites from crystal structure, it is difficult to know how many sites should be mutated and what types of mutations to choose, because enzyme activity is very sensitive to mutations, as well as the effects of epistasis^44^. Artificial intelligence (AI) models offer new approaches. Methods like ProteinMPNN, a method of protein inverse folding developed by David Baker et al.^42^, can better design stable protein sequences. To this end, we utilized ProteinMPNN to score the interface amino acids of both the dimeric and monomeric forms predicted by AlphaFold2 (AF2) (**Fig. S1**)^45^, obtaining two position-specific scoring matrices (PSSMs)^46^. Each matrix has 20 × M elements, where M is the length of the target sequence, and each element represents the MPNN score for a particular residue type at a given position. Subsequently, we conducted a correlation analysis of the MPNN scores of both the monomer and dimer (***EQ*** (**1**)). Sites with lower correlation may control PPI, as the MPNN scores of the dimer reflect the contribution of interface interactions, whereas those for the monomer do not (**Fig. 2a**).

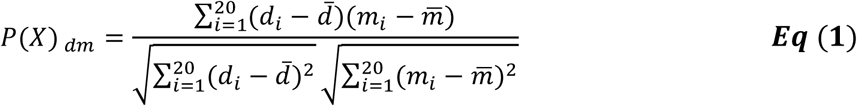

**Figure 2.**
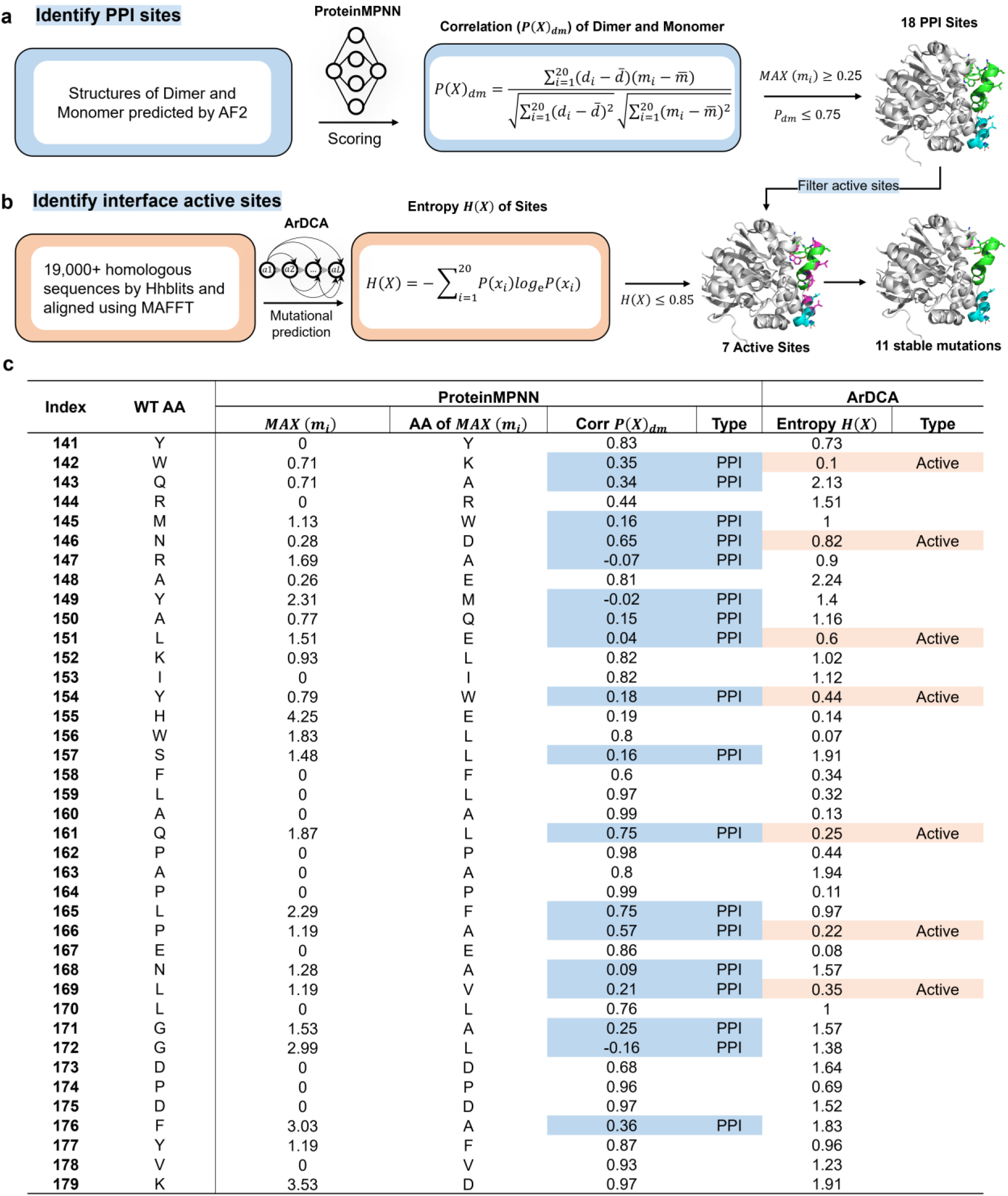
*De novo* design of the FAcD monomer using ProteinMPNN and ArDCA. **a**, The wild-type (WT) dimer and monomer structures predicted by AF2 were input into ProteinMPNN. After passing through the encoder and decoder modules, two PSSMs (di, mo) were obtained containing 302 sites. Sites with *MAX* (*m_i_)* ≥ 0.25 and correlation between the dimer and monomer ≤ 0.75 were identified as PPI sites. **b**, Over 19,000 homologous sequences were obtained by searching with HHblits, and a mutation matrix was also obtained using ArDCA. The entropy of each site was calculated separately. Sites with entropy ≤ 0.85 were considered conserved, and identified as potential active sites. These sites should not be mutated. **c**, MPNN scores, correlation and ArDCA entropy values of interface sites. From left to right, the wild-type amino acid position, amino acid type, amino acid with maximum monomer MPNN score, the maximum score, correlation between monomer and dimer scores, entropy of the site, and the type of each site.

Here X is the site, *d*_*i*_ represents the MPNN score of the site mutated to residue i in the dimer, *m*_*i*_ represents the MPNN score of the site in the monomer, and *d̅* and *m̅* are the averages of the 20 scores for the site.

To maintain the function of the monomer, we need to identify potential functional sites. In computational biology, it is commonly believed that more conserved sites are potential active sites. After searching a large number of homologous sequences and performing multiple sequence alignment (MSA), PSSM can be calculated. This PSSM also contains 20 × M elements, each of which is the log-likelihood of that particular residue type at that position, and the entropy of each site represents the degree of conservation at that position (**Eq (2)**). The lower the theoretical information entropy of a site, the more likely it is conserved and thus may affect activity^47^. However, the multiple sequences retrieved are limited and cannot fully represent the “natural” PSSM. Here, we employed HHblits^48^ to identify over 19,000 homologous sequences of FAcD and used ArDCA^43^ to predict a mutation matrix, represented as a PSSM. The PSSM predicted by ArDCA more accurately reflects natural distribution patterns compared to the one directly derived from the 19,000+ native sequences. This improvement can be attributed to the higher regularization coefficients in ArDCA, which better align mutation probabilities for each site more closely with natural distributions (**see Methods**). By excluding positions at the PPI interface that may affect activity, we can obtain a stable and active FAcD monomer (**Fig. 2b**).

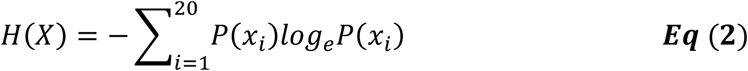

where X is the residue site, *P*(*x*_*i*_) represents the log-likelihood of X mutating into amino acid *x*_*i*_, and *x*_*i*_ represents 20 different amino acids.

The detailed MPNN scores for different structures are given in **Table S1-3**. The results demonstrated a strong Pearson correlation between the MPNN scores for each site of the monomer and dimer predicted by AF2, with an average correlation coefficient of 0.87. Moreover, the correlation coefficient between the monomer predicted by AF2 and the dimeric crystal structure (3R3U) was also high, with an average correlation coefficient of 0.9. Only a few sites had a lower correlation, and most of them were located in the dimer interface (Y141-K179), with a few located in the Loop B region (G249-T259) (**Fig. 2c and Fig. S2**). We then set the threshold to 0.75 to identify 18 different PPI sites and selected mutations with higher MPNN scores. Subsequently, using ArDCA with an entropy threshold of 0.85, we determined 7 potential active sites, with known active sites H155 and W156 having entropies below 0.2, which also proved that our strategy could identify potential active sites. Finally, after removing 7 active sites, we obtained a mutant with 11 stable mutations.

To test the effectiveness of the ProteinMPNN and ArDCA, we designed six mutants to test their stabilities and catalytic activities. **Table 1** summarizes the mutation sites of the six mutants. Among them, **Mu1-Mu3** are only considered for protein stability with ProteinMPNN and are expected to be inactive. **Mu1** has all 18 PPI sites replaced by amino acids with *Max* monomer scores. **Mu2** differs from **Mu1** only in restoring Y154 to test the effect of Y154W because this site is next to the active site H155. To further minimize interface mutations and avoid potential impacts on activity, **Mu3** targeted only the 9 sites with ProteinMPNN scores exceeding 1 that were located between residues M145–D175 and more than 5 Å away from the H155 center (excluding residues Y149-S157). In contrast, **Mu4**-**Mu6** incorporated refined interface mutations, while also considering active sites. The primary distinction between **Mu4** and **Mu5** lies in the L165F mutation. This position had a correlation coefficient of 0.75, which is right at the cutoff for inclusion (**Fig. 2c**). **Mu6** was derived from **Mu5** by removing the Y149M mutation to specifically investigate the importance of M149 in maintaining the cooperativity of the monomer.

**Table 1.**
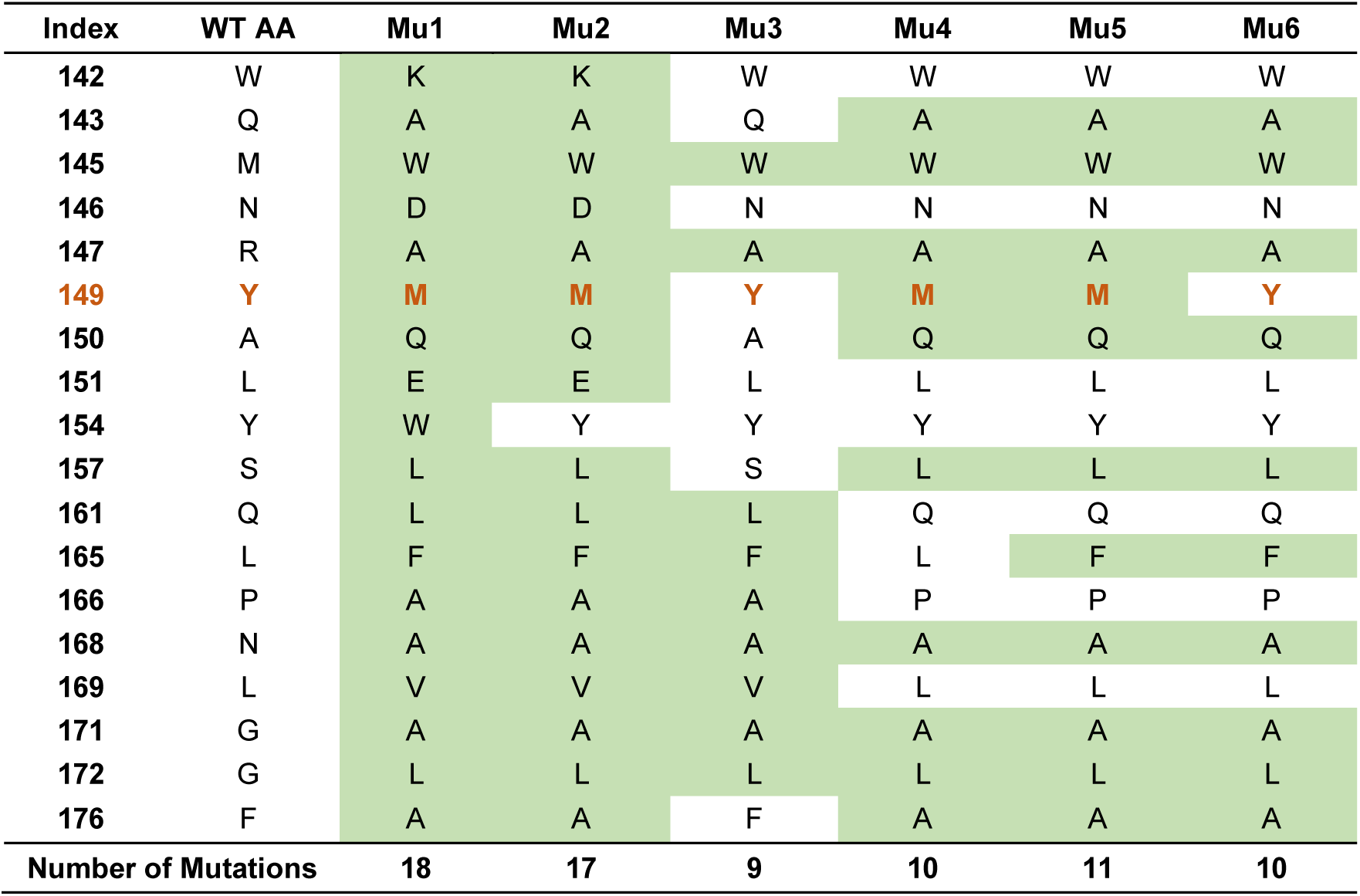
Mutation sites of six designed mutants. Mutants **Mu1**-**Mu3** only utilized the ProteinMPNN. **Mu1** involved all 18 PPI mutations; **Mu2** without Y154W mutations; **Mu3** targeted 9 sites with ProteinMPNN scores exceeding 1 that were located between residues M145–D175 and more than 5 Å away from the H155 center (excluding residues Y149-S157). **Mu5** removes the 7 active sites from **Mu1**; **Mu4** differs from **Mu5** without the L165F mutation; **Mu6** differs from **Mu5** without the Y149M mutation.

| Index | WT AA | Mu1 | Mu2 | Mu3 | Mu4 | Mu5 | Mu6 |
| --- | --- | --- | --- | --- | --- | --- | --- |
| 142 | W | K | K | W | W | W | W |
| 143 | Q | A | A | Q | A | A | A |
| 145 | M | W | W | W | W | W | W |
| 146 | N | D | D | N | N | N | N |
| 147 | R | A | A | A | A | A | A |
| 149 | Y | M | M | Y | M | M | Y |
| 150 | A | Q | Q | A | Q | Q | Q |
| 151 | L | E | E | L | L | L | L |
| 154 | Y | W | Y | Y | Y | Y | Y |
| 157 | S | L | L | S | L | L | L |
| 161 | Q | L | L | L | Q | Q | Q |
| 165 | L | F | F | F | L | F | F |
| 166 | P | A | A | A | P | P | P |
| 168 | N | A | A | A | A | A | A |
| 169 | L | V | V | V | L | L | L |
| 171 | G | A | A | A | A | A | A |
| 172 | G | L | L | L | L | L | L |
| 176 | F | A | A | F | A | A | A |
| Number of Mutations |  | 18 | 17 | 9 | 10 | 11 | 10 |

### Stability prediction and oligomeric form of mutants

To assess the stability of the mutants, we performed microsecond-scale MD simulations on both monomeric and dimeric forms. As shown in **Fig. 3a** and **3b**, all mutants exhibited strong stability in their monomeric forms. Specifically, for **Mu4** and **Mu5** monomers (**Mu4-Mo** and **Mu5-Mo**), the probability that the RMSD of Helix A relative to the AF2 structure was below 2 Å increased from 27% in **WT-Mo** to 75% and 66%, respectively. A similar trend was observed for Helix B, with RMSD below 2 Å rising from 7% in **WT-Mo** to 46% and 54%, respectively. In contrast, the dimeric interfaces of all mutants showed instability. For **Mu4** and **Mu5** dimers (**Mu4-Di** and **Mu5-Di**), the probability that the RMSD of Helix A was below 2 Å was only 7% and 2%, respectively, while for Helix B, the probabilities were 14% and 4%. These findings suggest that the mutants are stable as monomers but not as dimers, highlighting the utility of monomer-dimer correlations in identifying PPI sites and demonstrating the efficacy of ProteinMPNN in protein stability design.

**Figure 3.**
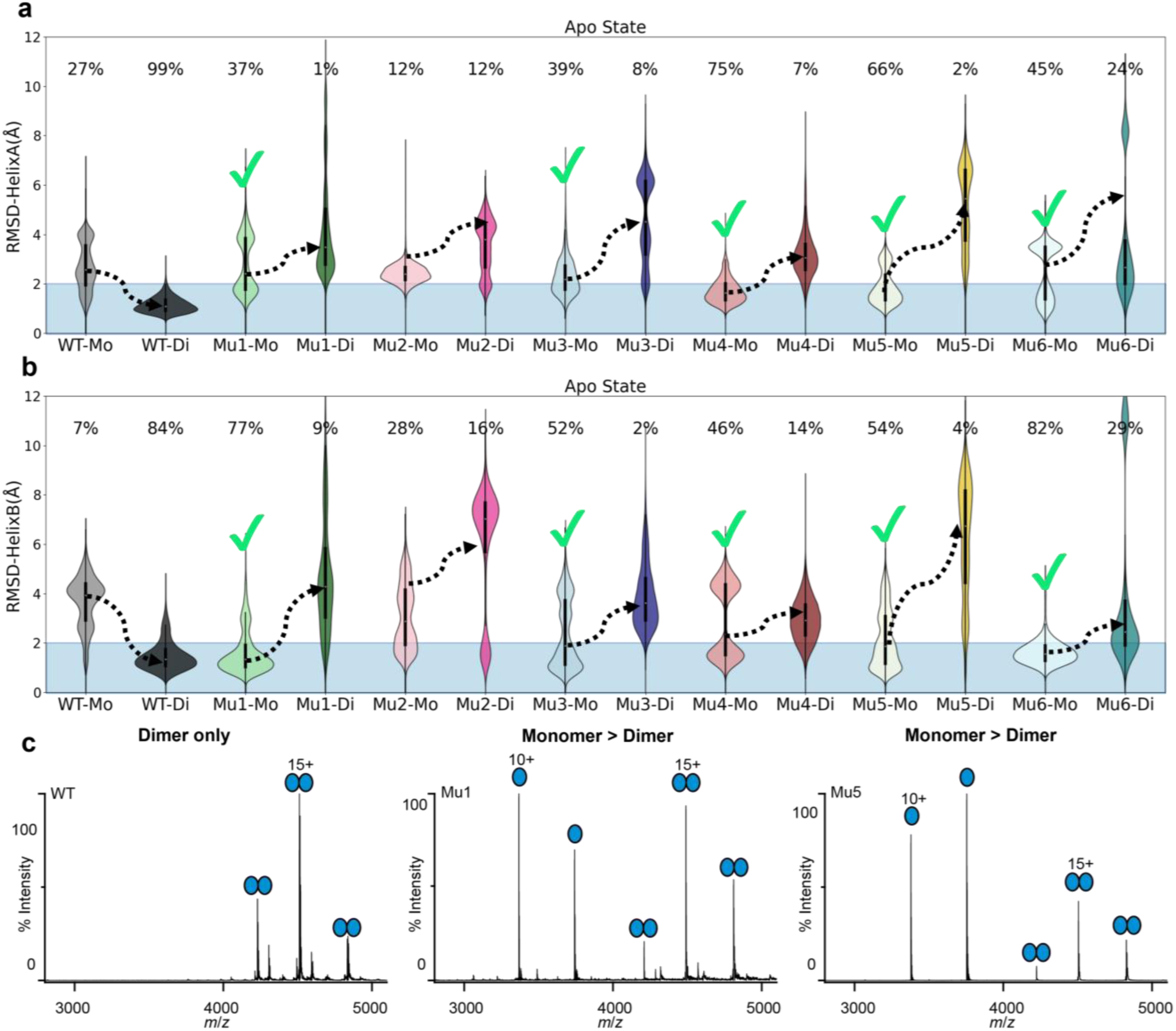
Stability of WT and mutants in MD simulations and oligomeric profiles by native MS. **a-b**, RMSD of Helix A and Helix B for the monomers and dimers of the wild-type and mutants (at 350 K for 3×1 μs). The percentages of RMSD below 2 Å for **WT** and mutants are marked. Key RMSD changes for both the monomeric and dimeric forms are highlighted with arrows, indicating significant stability deviations of the interface. **c**, Native MS spectra reveal the oligomeric profiles of **WT**, **Mu1**, and **Mu5**. The x-axis represents the mass-to-charge ratio (m/z), and the y-axis represents the relative intensity of the state. High, sharp peaks indicate a large number of charge states at a specific m/z value, while lower peaks indicate a low number of states. The wild-type exclusively forms a dimer, consistent with its stable dimeric configuration. In contrast, **Mu1** and **Mu5** exhibit significant shifts toward the monomeric state, which is the dominant signal in the spectra.

Following protein expression and purification, substantial amounts of mutant proteins were obtained, with expression levels exceeding those of the wild-type under identical induction conditions. To assess the oligomeric state of the mutants, **Mu1** (with 18 PPI mutations) and **Mu5** (with 7 active site mutations removed), we used mass spectrometry to analyze the wild-type and mutants at room temperature^49^. This approach allowed us to detect all oligomeric forms present in solution and to obtain the distribution of different charge states along with their relative intensities. As shown in **Fig. 3c**, the wild-type exists solely as a dimer (charge state 15+), with no monomer detected. In contrast, **Mu1** and **Mu5** both exhibited monomer and dimer peaks, with the monomer being the predominant spectral signal (charge state 10+). These findings are in qualitative agreement with our predictions.

After combining the trajectories of **WT** and mutants, PCA analysis was performed on the non-hydrogen atoms of the pocket residues (D110, H155, H280). The results (**Fig. S3a**) revealed similar conformational distributions for **Mu4-Mo**, **Mu5-Mo**, **Mu6-Mo**, and **WT-Di**, suggesting they are more likely to exist in a monomeric form while retaining catalytic activity. Although **Mu1-Mu3** are stable in their monomeric forms, their pocket conformations deviate from that of **WT-Di**. Structural analysis of the mutants indicates that 7 active residues are critical for maintaining the catalytic conformations of Helix A and Helix B. Mutations at these positions in **Mu1-Mu3** disrupt the interaction network, preventing H155 from participating in catalysis, implying that these mutants are inactive. In contrast, the active sites in **Mu4-Mu6** remain intact, and H155 retains its catalytic function within the pocket (**Fig. S3b**).

### Activity prediction and kinetic experiments of mutants

To further investigate the activity of the mutants, we conducted detailed mechanistic studies. In this study, we employed α-Fluorophenylacetic acid (**1aa**), which exhibits a *k*_cat_ of 676.3 min^-1^, as a substrate for mechanistic investigation^34^. This substrate demonstrates significantly higher activity compared to the natural substrate fluoroacetate (**FAc**)^32^, which has a *k*_cat_ of 1.84 min^−1^. Substrate **1aa** undergoes defluorination, nucleophilic attack by water, and subsequent C-O bond cleavage to yield the final product, with the defluorination and nucleophilic attack steps being particularly critical^31^. Accordingly, we defined the pre-reaction states (PRS)^50^ of defluorination and nucleophilic attack based on the distances and angles and calculated the proportion of active conformations in trajectories (**Fig. 4a-c**). As shown in **Fig. 4d**, while the interfaces of **Mu1-Mo**, **Mu2-Mo**, and **Mu3-Mo** are stable during defluorination and nucleophilic attack (**Fig. S4**), their catalytic environments may be disrupted, preventing stabilization in the PRS conformation, with a PRS occupancy of 0.0% during the defluorination step. Structural analysis showed that the H155 side chain swings out from the pocket to the protein surface, leading to a loss of activity (**Fig. S5-6**). In contrast, **Mu4-Mo** and **Mu5-Mo** exhibit defluorination PRS ratios of 5.4% and 4.6%, respectively, similar to **WT-Di** (3.7%), indicating that **Mu4-Mo** and **Mu5-Mo** are potential active monomers. **Mu6-Mo** retains some defluorination activity, but with the loss of Y149M, its active conformations drop to 2.1%, lower than **WT-Di** (**Fig. 4d**).

**Figure 4.**
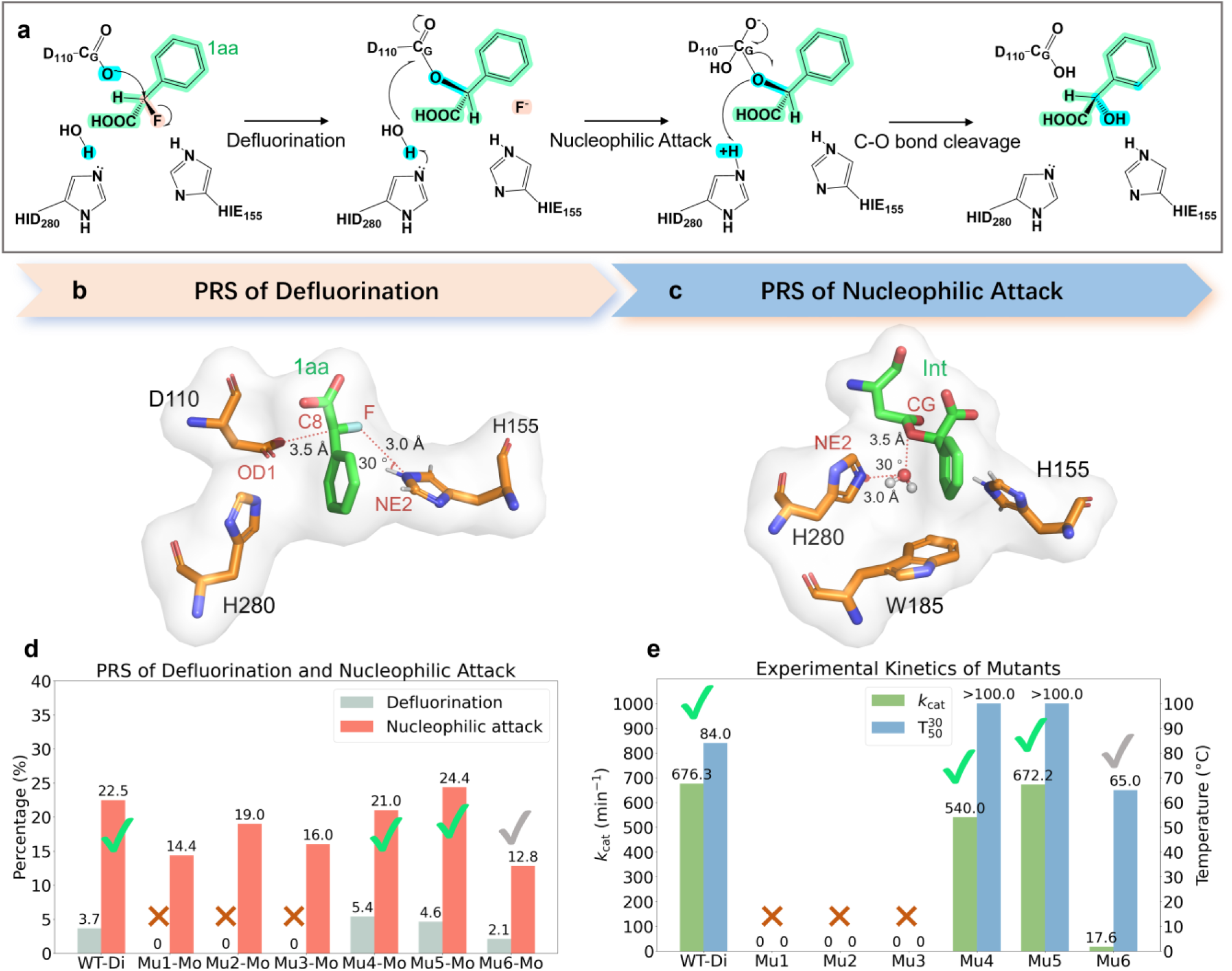
Definition of the Pre-reaction states for defluorination and nucleophilic attack and results of kinetics experiments. **a**, The mechanism by which FAcD catalyzes the conversion of substrate **1aa** to the product. **b-c**, The active conformations defined by PRS for substrate defluorination and nucleophilic attack. **d**, The ratio of active conformations in **WT** and mutants in all trajectories during the defluorination and nucleophilic attack. The molecular dynamics simulations for the defluorination step were performed for 3 × 200 ns, while the nucleophilic attack simulations were carried out for 3 × 1 µs. **e**, The *k*_cat_ values obtained for **WT** and mutants experimentally, along with their T_50_^30^ (the temperature at which activity is reduced by half after 30 minutes of incubation).

For the nucleophilic attack step, it was similar to those of defluorination, with the probabilities of active conformations for **Mu1-Mo**, **Mu2-Mo**, and **Mu3-Mo** being 14.4%, 19.0%, and 16.0%, respectively, lower than **WT-Di** (22.5%). While **Mu4-Mo** and **Mu5-Mo** also had the ability to perform nucleophilic attack similar to **WT-Di**, with active conformations ratios of 21.0% and 24.4%, respectively. Inversely, **Mu6-Mo**’s active conformations were only 12.8%, significantly lower than that of the **WT-Di**, possibly due to the loss of Y149M, which will be discussed in more details in the next section. In summary, we speculate that **Mu4** and **Mu5** are potential active monomers, while **Mu6** can also catalyze, but with lower efficiency. **Mu1-Mu3** are limited by the defluorination and nucleophilic attack steps and are inactive monomers.

Then we conducted a detailed kinetic validation of **WT** and mutants to further validate our calculation. As depicted in **Fig. 4e**, both experimental and computational results were consistent, with **Mu1**, **Mu2**, and **Mu3** showing no activity (*k*_cat_ = 0), while **Mu4** and **Mu5** exhibited catalytic activity. **Mu4** had a *k*_cat_ of 540.0 min^-1^, and **Mu5** had a *k*_cat_ of 672.2 min^-1^, values comparable to that of **WT-Di** (676.3 min^-1^)^34^. The L165F mutation in **Mu5** resulted in a 1.24-fold increase in activity over **Mu4**, further confirming the accuracy of ArDCA in predicting potential active sites. In contrast, entropy values directly calculated from 19,000+ native homologous sequences showed poor dispersion, making it challenging to identify active sites. Even with a cutoff value of 1.6, nine potential active sites were identified, with L165 being identified as a key active site that should not be mutated (**Fig. S7**). Moreover, the Y149M mutation in **Mu5** significantly enhanced its activity nearly 40-fold compared to **Mu6**, highlighting the critical role of Y149M in boosting catalytic efficiency. MD simulations, however, failed to identify Y149 as a critical PPI site based on contact frequency analysis by GetContacts^51^, with a contact frequency cutoff of 0.6 (**Table S4**). These findings underscore the efficacy of combining ProteinMPNN and ArDCA for the accurate and rapid identification of both PPI and active sites, without the need for extensive structural analysis.

Stability analysis showed that after 30 minutes at 84 °C, the activity of **WT-Di** decreased by half, likely due to dissociation into a non-active monomer (**Fig. S8**). In contrast, **Mu4** and **Mu5** retained near-full activity after 30 minutes at 100 °C (**Fig. 4e**), suggesting that these mutants remain stable as monomers and are resistant to high-temperature dissociation. This enhanced thermal stability and higher expression levels of the monomers not only simplify industrial purification but also reduce production costs by enabling high-temperature purification and direct yield of concentrated enzyme protein.

### Y149M Restores Cooperativity in Mu5-Mo from WT-Di

The marked difference in catalytic activity between **Mu5** and **Mu6** is striking. The Y149M mutation in **Mu5** leads to a nearly 40-fold increase in catalytic activity compared to **Mu6**, which lacks this mutation. Our preliminary investigations demonstrated the existence of inter-subunit cooperativity within the FAcD dimer^36^. As shown in **Fig. 5a**, the substrate binding to the non-catalytic subunit forms a hydrogen bond with Y141’ (all residues of the **NC** subunit are annotated with superscript primes) and induces a pronounced proton chain at the interface: Y141-H₂O-S157-H₂O-S157’. Due to the *π*-*π* interactions between Y141 and Y149, the opening of Y141 on the catalytic subunit is accompanied by the displacement of Y149, which moves away from Loop B without disrupting it. The stable Loop B restricts W185 in an open conformation, facilitating water molecule entry for catalysis^36^. Although **Mu5**, as a monomer, the small side-chain amino acid M149 is unlikely to disrupt Loop B. Therefore, **Mu5** may preserve the cooperative efficiency within the monomer. In contrast, **Mu6**, containing Y149, may destabilize Loop B. This disruption would prevent loop B from maintaining W185 in its open conformation, hindering water molecule entry and adversely affecting catalytic activity.

**Figure 5.**
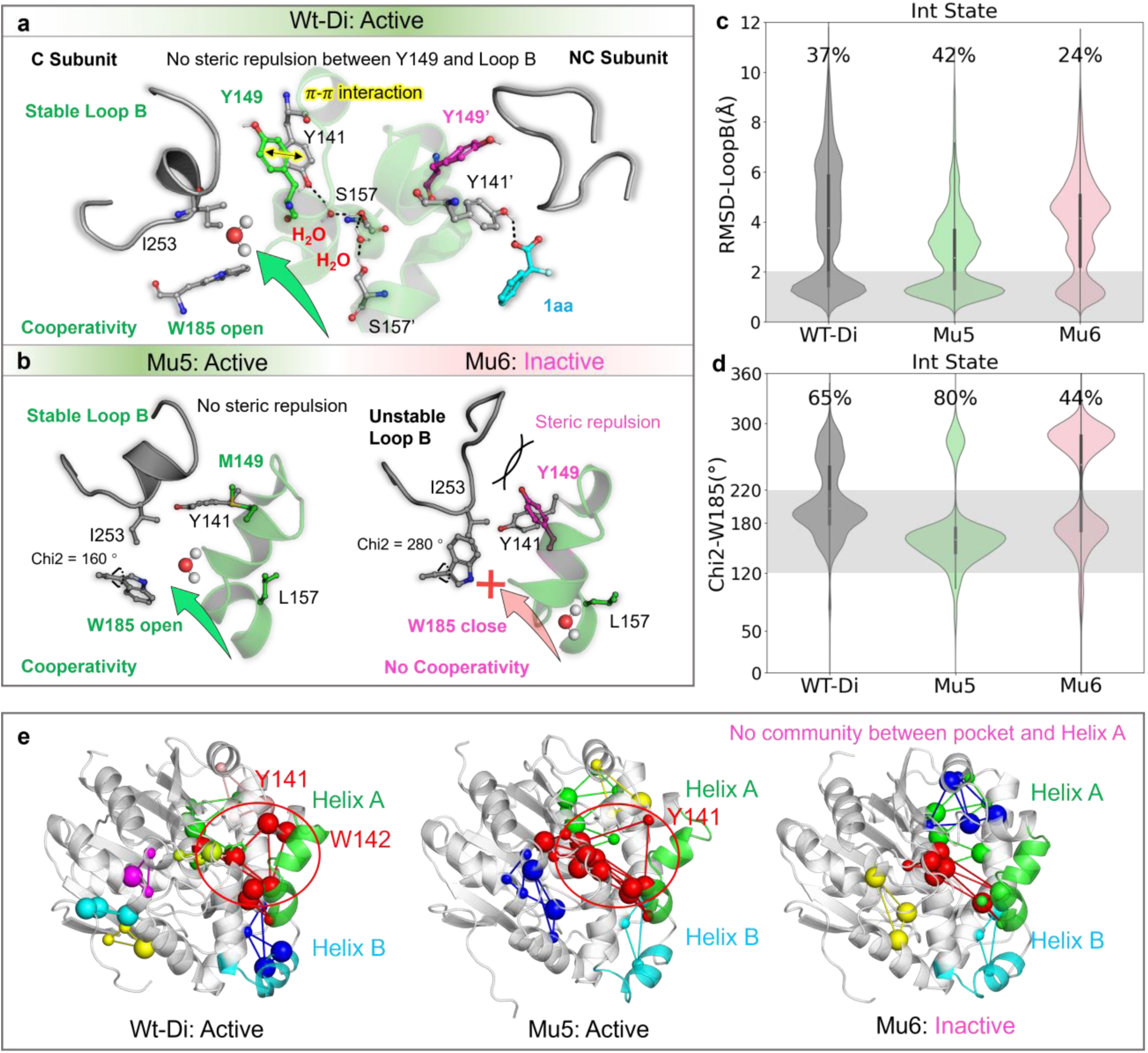
Y149M Enhances Catalytic Efficiency in Mu5. **a,** The key proton chain responsible for cooperative catalysis in **WT-Di**. The substrate in the **NC** subunit induces the formation of this proton chain at the interface, causing Y141 and Y149 in the **C** subunit to open, reducing steric hindrance on Loop B. Stable Loop B locks W185 in an open state, facilitating water molecule entry for catalysis. **b,** Optimal conformations of **Mu5-Mo** and **Mu6-Mo** obtained via Dpeaks clustering. In **Mu5-Mo**, M149 does not disrupt Loop B, and W185 remains open. In contrast, Y149 in **Mu6-Mo** disrupts Loop B, and W185 adopts a closed conformation. **c-d,** RMSD distribution of Loop B and Chi2 angle distribution of W185 for **WT-Di**, **Mu5-Mo**, and **Mu6- Mo**. **e,** Community analysis based on webPSN. All 3 × 1 µs trajectories were used for Protein Structure Network (PSN) generation and community analysis.

To further elucidate the importance of Y149M, MD simulations were conducted for **Mu5-Mo** (M149) and **Mu6-Mo** (Y149) and the pocket amino acids were clustered by Dpeaks^52^ to obtain representative conformations. As shown in **Fig. 5b**, in **Mu5**, M149 as a small side-chain amino acid does not interfere with Loop B. Stable Loop B restricts W185 to close upward, adopting an open conformation favorable for water molecule entry. However, if M149 is mutated to Y in **Mu6**, Y149 significantly disrupts Loop B. As a result, W185 easily rotates upward to close, thus preventing water molecules from entering. Specifically, compared to **WT-Di**, the proportion of Loop B with an RMSD below 2 Å in **Mu5** increased from 37% to 42%, whereas in **Mu6**, this value was only 24%. Additionally, the probability of W185 adopting an open conformation in **Mu5** was also comparable to that in **WT-Di**, increasing from 65% to 80%. While in **Mu6**, this probability remained at only 44%. (**Fig. 5c, d**). Kinetic results also show that the activity of **Mu6** is significantly reduced by nearly 40-fold, from 672.2 min^-1^ to 17.6 min^-1^.

webPSN^53^ can identify communities of closely related sites, playing a key role in the study of protein allosteric regulation. These community sites are often motion-related and collectively maintain the stability of the community. In our community analysis of nucleophilic attack conformations, we found that the pockets of **WT-Di** and **Mu5-Mo** can form communities with Helix A, while the pockets of **Mu6-Mo** cannot (**Fig. 5e**). This suggests that the interface of **Mu5** stabilizes the pocket for catalysis, whereas the Y149 mutation in **Mu6** creates steric hindrance with Loop B, reducing the interface-pocket interaction and decreasing catalytic activity. These results validate that the Y149M mutation enhances cooperativity in monomer and demonstrate that our strategy effectively ensures the stability, activity, and cooperative effects of the FAcD monomer.

## Discussion

Converting dimers or oligomers into active monomers is a significant challenge, as it requires not only ensuring monomer stability but also preserving enzyme activity, especially, when the dimer interfaces overlap with the catalytic center. Advances in deep neural networks have revolutionized generative protein design^54^, making it possible to design *de novo* proteins with specified folds^42, 55^. ProteinMPNN, an inverse folding model, learns the relationship between structure and sequence to generate protein sequences that stably fold. On natural protein backbones, ProteinMPNN achieves a sequence recovery rate of 52.4%^42^. Using this model, our designed FAcD monomers demonstrated enhanced stability.

Generative protein sequence models can capture the relationship between natural sequence distributions and potential functions^43, 56, 57^. ArDCA, for instance, utilizes maximum-likelihood inference on multiple sequence alignments of homologous proteins to train an autoregressive model for sequence generation and mutation prediction^43^. In mutation prediction, applying larger regularization terms biases the predicted distribution toward “natural” patterns, effectively capturing the true conservation at each site. Using ArDCA, we identified 7 accurate active sites, resulting in the successful design of active monomers **Mu4-Mu6**. Incorporating the Y149M mutation transferred the cooperative effect of the dimer to the monomer, yielding monomeric activity comparable to that of the wild-type dimer. Accurately identifying potential active sites and cooperative catalytic sites is therefore crucial, particularly the latter. A single-site mutation can dramatically enhance catalytic activity, increasing turnover rates from 17.6 min^-1^ to 672.2 min^-1^. Machine learning models capable of precisely predicting cooperative catalytic sites could revolutionize enzyme design by overcoming the time and technical limitations of traditional directed evolution approaches, which often target pocket residues.

By integrating ProteinMPNN and ArDCA, we achieved the *de novo* design of the FAcD monomer without extensive manual consideration of enzymatic properties. This approach effectively identified PPI sites and key residues affecting activity. Notably, mutant **Mu5** exhibited activity similar to the wild-type dimer but with enhanced stability (T_50_^30^ > 100 ℃). The monomer design strategy has many important applications. (1) It enables the control of the oligomeric form of enzymes while maintaining high activity. This process is not limited to converting dimers to monomers, but also applicable to transforming any higher-order oligomers to lower-order ones, provided the structure of the protein is available. In some cases, the higher-order oligomeric forms of enzyme proteins, rather than just dimers, often lack activity^58, 59^. Stabilizing the active form of a protein is crucial; (2) It aids in the preservation of proteins. Proteins can lose activity due to abnormal oligomerization after long-term storage, but monomers will not; (3) This strategy can be used to reduce self-inhibition in functional proteins; (4) if the monomeric form of an enzyme can achieve the stability and activity of dimers/higher-order oligomers, the immobilized carrier can adsorb more enzymes due to the low steric effect of the monomer; (5) Additionally, stable monomers can act as scaffolds for artificial enzymes^60^, such as the TIM barrel used in designing retro-aldolase RA95^61, 62^, as well as for creating new artificial metalloenzymes for the enantioselective epoxidation of cellulose^63^.

Recently, there have been many tools for predicting PPI sites^64–66^, as well as similar multimer prediction tools like AF2-multimer^67^, but these tools need the specification of structures or oligomeric forms, which cannot truly predict the oligomeric state from a sequence. We aim to focus on the oligomeric form of enzymes while engineering enzymes, and it would be particularly important to develop sequence-based predictors for oligomeric forms. This is also important on the basis of existing *k*_cat_ predictors^68–70^, when the enzyme oligomeric state is considered. This work underscores the importance of controlling oligomeric states in enzyme biotechnology, particularly for immobilization applications, and highlights its potential for designing functional proteins. By combining computational advances with experimental validation, we aim to push the boundaries of enzyme engineering and protein design.

## Methods

### Protein preparation

Genes encoding the Rpa1163 gene (1-302) were cloned into a modified pRSF-Duet vector, preceded by a His6-SUMO tag. The fusion proteins were over-expressed in *E. coli* BL21(DE3) cells, which were induced by the addition of IPTG at an OD600 of 0.8 and then grown at 16 ℃ for 12-16 hrs. The cells were harvested and lysed in a buffer containing 25 mM Tris-HCl (pH 8.0), 1 M NaCl, 25 mM imidazole, 0.5 mM β-ME and 1 mM PMSF. The fusion proteins were first purified using a nickel column. The protein was washed using washing buffer with 25 mM imidazole, 1M NaCl, 25 mM Tris-HCl (pH 8.0). Then the protein was eluted with elution buffer (250 mM imidazole, 1M NaCl, 25 mM Tris-HCl and pH 8.0). The His6-SUMO tag was cleaved off by ULP1 and the Rpa1163 (1-302) protein was fractioned using Nickel column again, and size exclusion column (SuperdexTM 75 Increase 10/300 GL, GE Healthcare) pre-equilibrated with a buffer containing 50 mM Tris-H_2_SO4 (pH 8.5), 150 mM NaCl and 3 mM DTT, finally buffer-exchanged into 50 mM Tris- H_2_SO4 pH 8.5.

### Native Mass Spectrometry

Proteins were buffer-exchanged into 0.2 M ammonium acetate (pH 7) and analyzed by a Q-Exactive Plus Hybrid Quadrupole-Orbitrap Mass Spectrometer^49^. Spectra were acquired based on an established protocol with the collision voltage set at 20 V^71^. Mass and charge states were determined using an Excel script kindly provided by the Benesch Laboratory, University of Oxford^72^.

### Kinetic of WT and mutants

To determine the kinetic parameters of **WT** and mutants with *α*-fluorophenylacetic acid as the substrate, a reaction mixture containing *α*-fluorophenylacetic acid (2-40 mM), 1.0 μM mutant in a total volume of 300 μL Tris-H_2_SO_4_ buffer (50 mM, pH = 7.0) was incubated at 60 °C and 1200 RPM for 24 h. The reaction mixture was quenched by adding 30 μL concentrated HCl, followed by extraction with 500 μL EtOAc. Subsequently, the mixture was centrifuged at 1200 RPM for 3 min, and 400 μL organic supernatants were transferred to a 2 mL centrifuge tube. The methylation was performed by adding 240 μL of methanol and 30 μL of (Trimethylsilyl) diazomethane solution, and then incubated at 30 °C, 800 RPM for 30 min. The consumption of the substrate was determined by comparing the peak area of two enantiomers in Gas chromatography. Data fitting was performed using Origin 2022b.

The T ^30^ of **WT** and mutants were tested at different temperatures for 30 min, and the remaining conditions were the same as the above kinetic tests.

### Homologous sequence search and ArDCA

Multiple homologous sequence searches were conducted using HHblits^48^, with the protein sequence (Uniprot_id: Q6NAM1) serving as the query. The search was meticulously conducted, setting the coverage threshold (-mact) to 0.35, where databases UniRef30_2020_06 and bfd_metaclust_clu_complete_id30_c90_final_seq.sorted_opt were employed, and E-value was set to 1e^-10^, ensuring high-quality results. Subsequently, hhfilter in HHsuite package^73^ was used to filter these sequences, retaining sequences with a maximal sequence identity of 99% (--id 99) and ensuring a minimal coverage with the query of 75% (--cov 75), resulting in 19,377 homologous sequences. Finally, MAFFT^74^ was used for multiple sequence alignment, employing the FFT-NS-2 method and setting the maximum iteration count to 3 (--retree 3) to optimize the alignment output.

After multiple sequence alignment of the 19,377 sequences, the PSSM can be directly calculated, and then the entropy of each site is obtained according to ***EQ*** (**1**).

Mutation prediction using ArDCA^43^ necessitates relatively large regularization parameters (*λ*_*J*_ = 10^−2^, *λ*_*H*_ = 10^−4^). Smaller regularization tends to generate sequences that closely resemble the input sequences, but for mutation effects prediction, a larger regularization is required, as the model needs to exhibit stronger generalization capabilities from the limited input. For a detailed tutorial, please refer to: https://github.com/pagnani/ArDCA.jl/blob/master/julia-notebook/tutorial.ipynb. Then the mutation probabilities of 20 amino acids at each site are obtained, and entropy calculation is performed according to ***EQ*** (**1**).

### ProteinMPNN

After predicting mutation effects using ProteinMPNN^42^, a mutation probability matrix is obtained for each residue position. For a detailed tutorial, please refer to the ProteinMPNN GitHub repository: https://github.com/dauparas/ProteinMPNN.

### MD simulations and analysis

We used the prediction model of AlphaFold2 as the initial structure for subsequent simulations^45^. The protonation state of the model was modified using the H++ server^75^. The protonation states of H155 and H280 were strictly set according to the previous mechanism research to avoid significant fluctuations in the pocket amino acids^31^.

The parameters for standard amino acid residues in the enzyme were taken from RSFF2C, an adapted version of the ff14SB force field developed by our group^76–78^. Gaff2 force field parameters were selected for small molecules and covalent ester intermediates^79, 80^. ach system was solvated in a TIP3P water model in an octahedral box. The charges of the small molecule 1aa, its ester intermediate, and the product were calculated by RESP2 using Multiwfn^81, 82^. Gaussian 16 was used for geometric optimization (B3LYP-D3(BJ)/def2-SVP) and single point calculation (B3LYP-D3(BJ)/def2-TZVP)^83^.

All systems are treated with periodic boundaries, and the closest distance between the protein surface and the box is 9 Å. Long-range electrostatic interactions were calculated using the Particle Grid Ewald method with a non-bonding cutoff of 9 Å. The temperature is controlled by the Langevin thermostat with the collision frequency set to 2 ps^−1^ in all simulations. An integration time step of 2 fs was used, and the SHAKE algorithm was employed to constrain bonds involving hydrogen atoms. All simulations were carried out under the NVT system.

First, the initial structure needs to be pre-balanced, and the Cα atoms are restricted to eliminate unreasonable interactions. Subsequently, the system was stabilized from 0K to 350 K within 40 ps and held at this temperature for 100 ps. Finally, high-temperature MD simulations were run at 350 K for up to 1 μs. Three independent simulations were run for each system, so a total of 3×1 μs MD simulations were performed. Simulations of all systems were performed with the Amber20 software using the GPU-accelerated pmemd module^79^, saving structures every 40 ps.

The RMSD of Helix A, Helix B, and Loop B are calculated with the predicted structure of AF2 as the reference conformation in all trajectories. All trajectories were subjected to Dpeaks cluster analysis^52^. This study focuses on pocket amino acids, so the non-hydrogen atoms (D110, H155, H280) of pocket amino acids are selected as the research objects. The epsilon between conformations was selected as 1, and the most dominant conformation in the thermodynamic diagram was selected among the major categories with the highest proportion. All figures were drawn with the help of Pymol^84^.

### Restricted simulation and pre-reaction state analysis

When exploring the process of substrate defluorination, the substrate needs to be constrained to an active conformation in the catalytic pocket to enhance PRS sampling: the distance between F of 1aa and NE2 of H155 and NE1 of W156 is kept at 2.8–3.2 Å, while the distance between C8 of 1aa and the O of D110 remains at 3.3–3.7 Å. The force constant is 10 kcal / (mol Å^2^). The distance of the pre-reaction state is defined as the distance between F of 1aa and NE2 of H155 and NE1 of W156 is less than 3.0 Å, while the distance between O atom of D110 and C8 of 1aa is less than 3.5 Å. Finally, a high-temperature MD simulation was run at 350K for 200 ns. Three independent simulations were run for each system, so a total of 3 × 200 ns MD simulations were performed.

When exploring the process of nucleophilic attack, the distance of the pre-reaction state is defined as the distance between NE2 of H280 and H of water molecule is less than 3.0 Å, while the distance between the O of water molecule and CG of Int is less than 3.5 Å.

The average number of water molecules in the three trajectories is calculated by the following formula ***EQ*** (**3**):

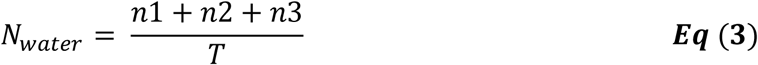

Among them, *N_water_*is the average number of the three trajectories, and the values of n1, n2, and n3 are taken as 0 or 1, respectively representing the number of active conformations contained in the three trajectories at time T, and the value of T is from 0 ns to 1000 ns.

## Acknowledgments

This work was funded by the National Key Research and Development Program of China (No. 2023YFA1506500, 2023YFA0914100/2023YFA0914102 to J. W.). The Key-Area Research and Development Program of Guangdong Provinces (2020B0101350001). Shenzhen Fundamental Research Program (No. GXWD20201231165807007- 20200812124825001) and the National Natural Science Foundation of China (No. 22403067, 22477110 to J. W.). This work was supported by Shenzhen Bay Laboratory Supercomputing Center.

