## Supplemental figures and tables for "A *De Novo* Design Strategy to Convert FAcD from Dimer to Active Monomer"

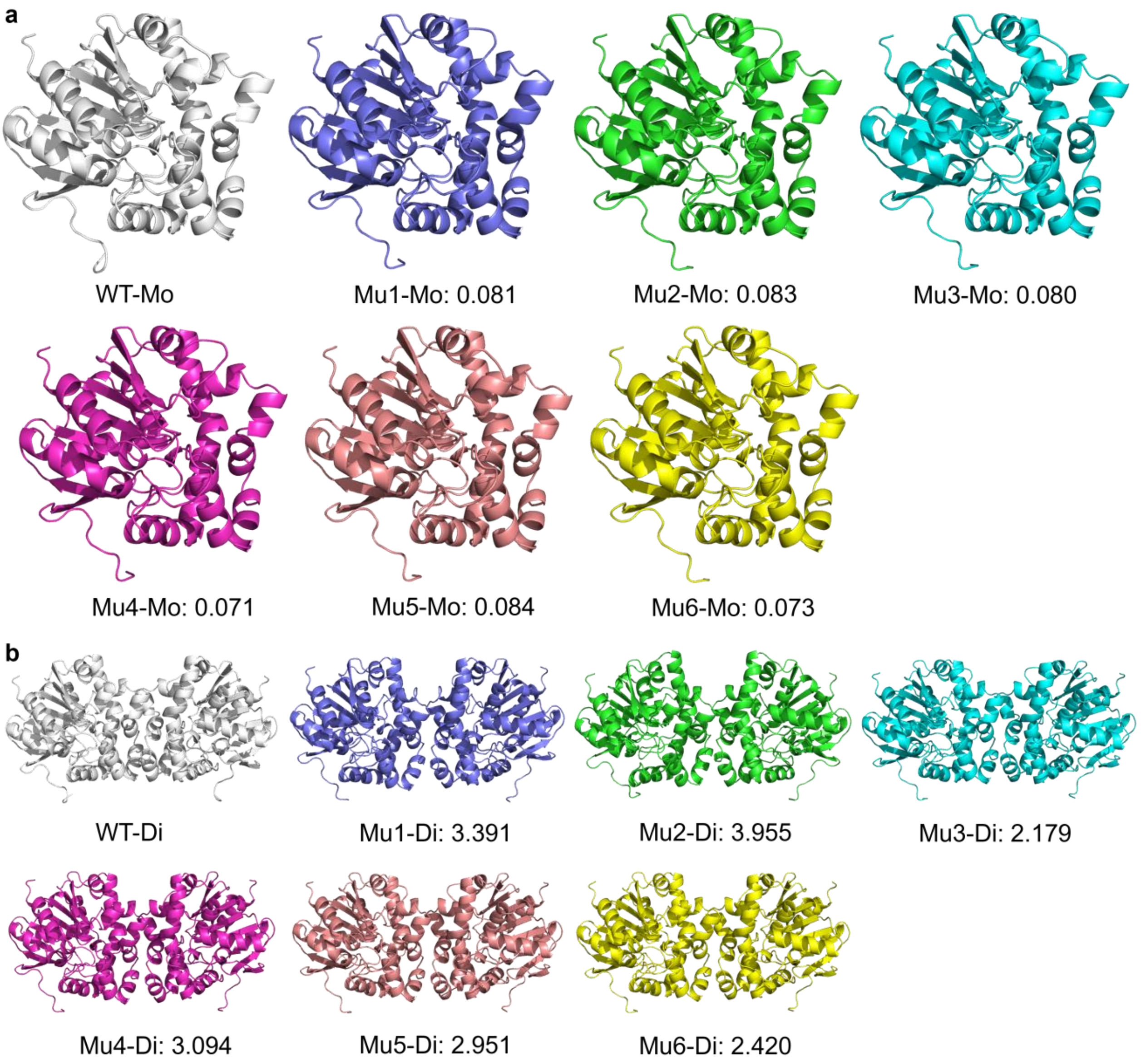

**Figure S1. Structure of mutants predicted by AF2.** The RMSD of all Cα of mutants compared with **WT-Mo** (**a**) and **WT-Di** (**b**) are labeled.

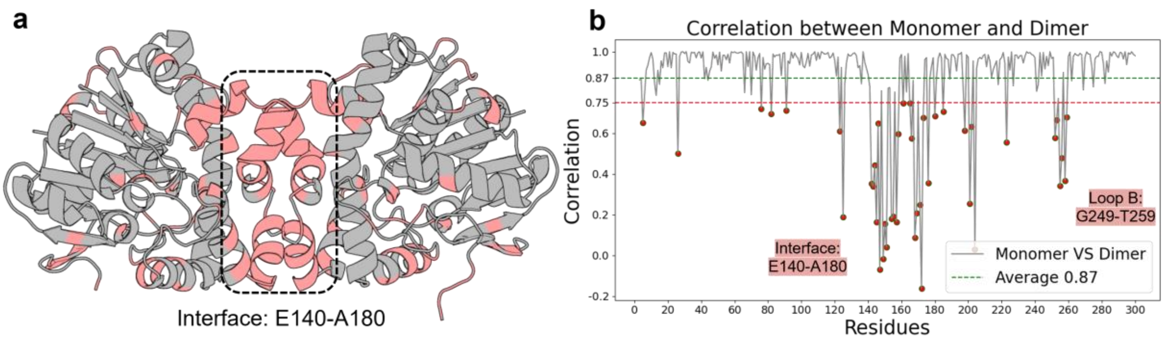

**Figure S2. PPI sites with low correlation at the interface. a,** Sites with correlation between the dimer and monomer < 0.87 (the average value) were marked in red on the AF2 dimer structure. **b,** Sites with a correlation < 0.75 were marked.

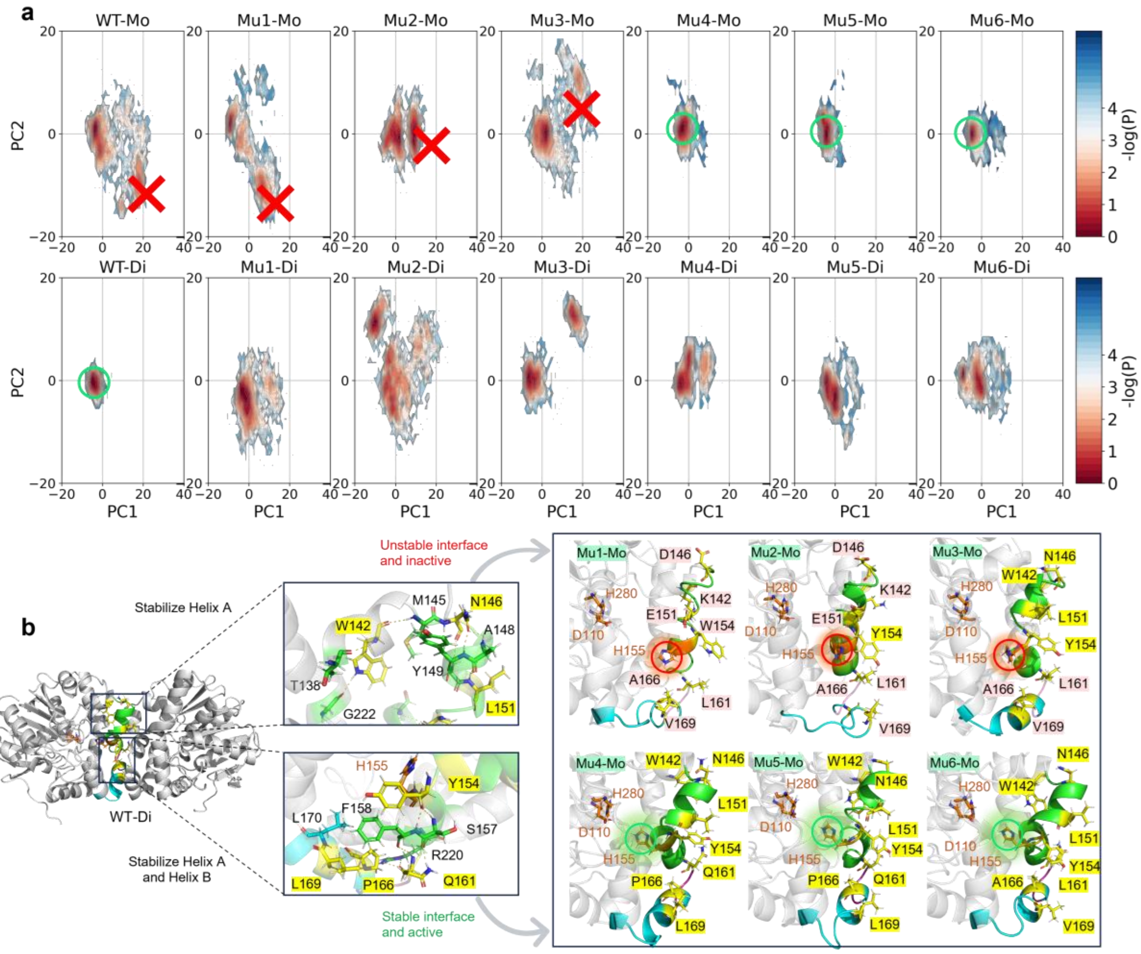

**Figure S3. PCA results of the pocket amino acids of the wild type and mutants. a,** After merging trajectories in different states, PCA was performed on the pocket amino acids (D110, H155, H280, non-hydrogen atoms) of all conformations of 3×1us. **b,** Representative conformations of wild type and mutants obtained by Dpeaks clustering.

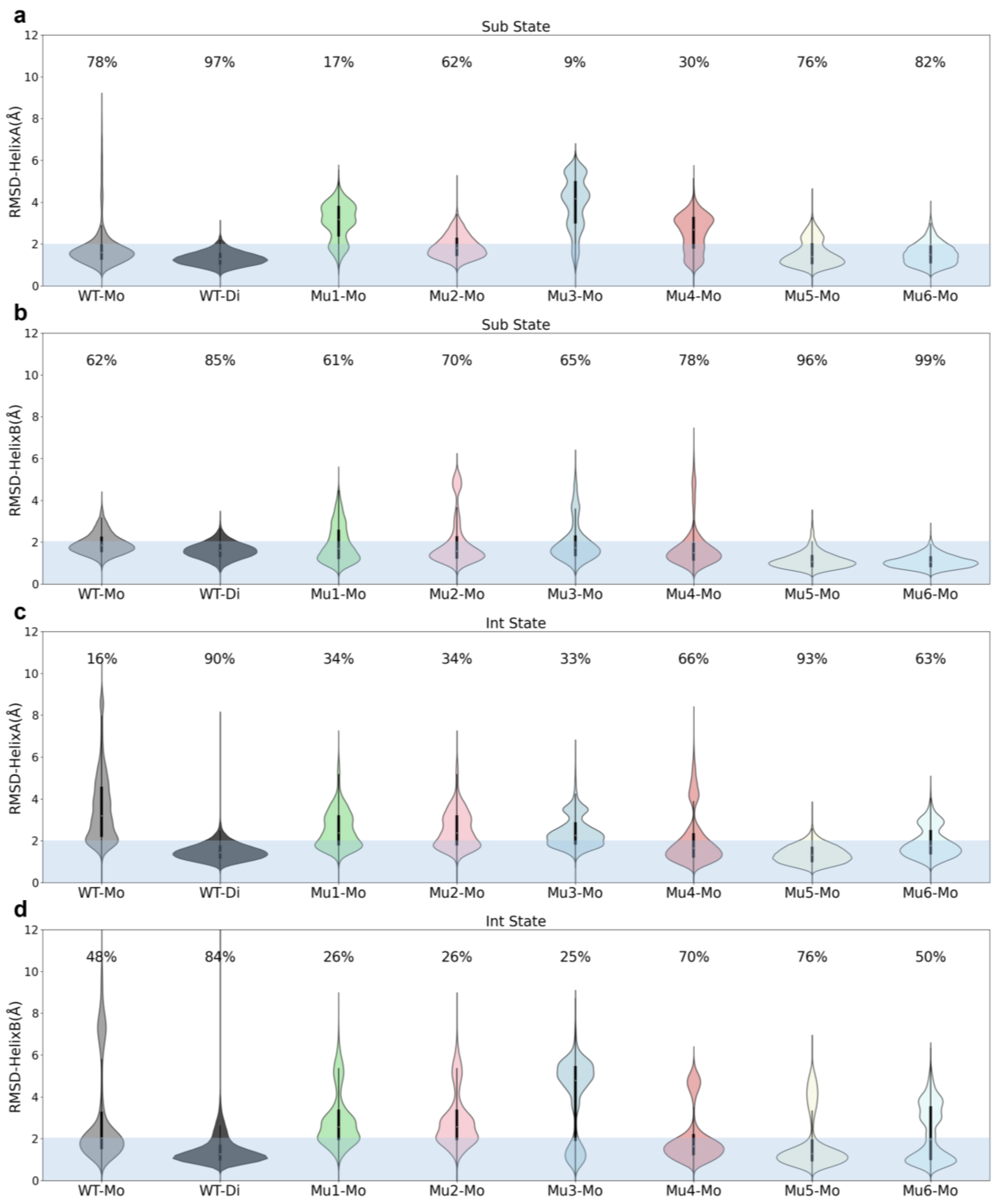

**Figure S4. RMSD of Helix A and Helix B of WT and mutants in the Sub and Int states. a-b**, RMSD of Helix A and Helix B of WT and mutants in Sub state. **c-d**, RMSD of Helix A and Helix B of WT and mutants in Int state.

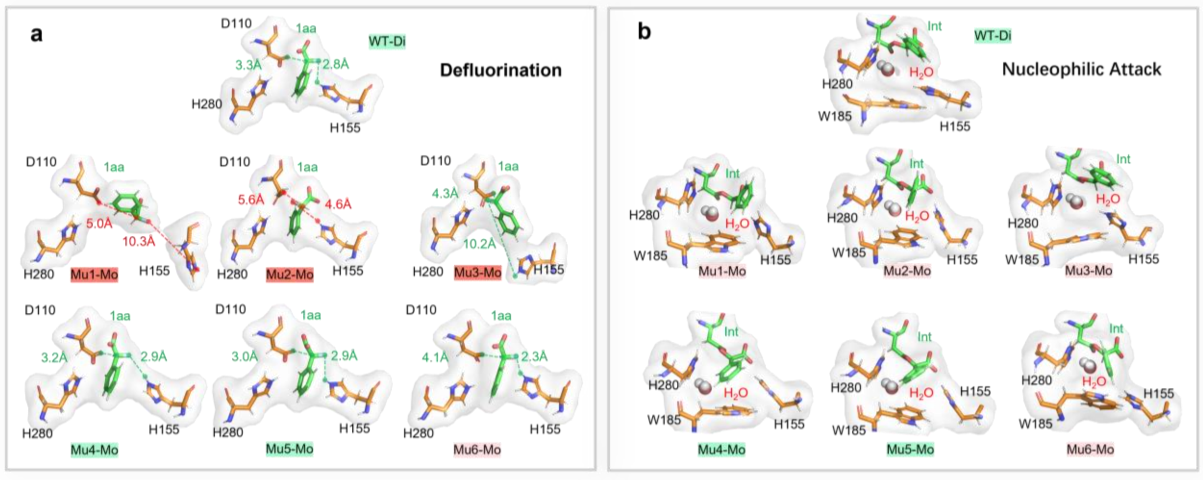

**Figure S5. Representative conformations of WT and mutants during defluorination and nucleophilic attack. a,** Representative conformations of wild-type and mutants for defluorination were obtained by clustering pocket amino acids (D110, H155, H280, non-hydrogen atoms) with the Dpeaks. **b,** Representative conformations of wild-type and mutants for nucleophilic attack.

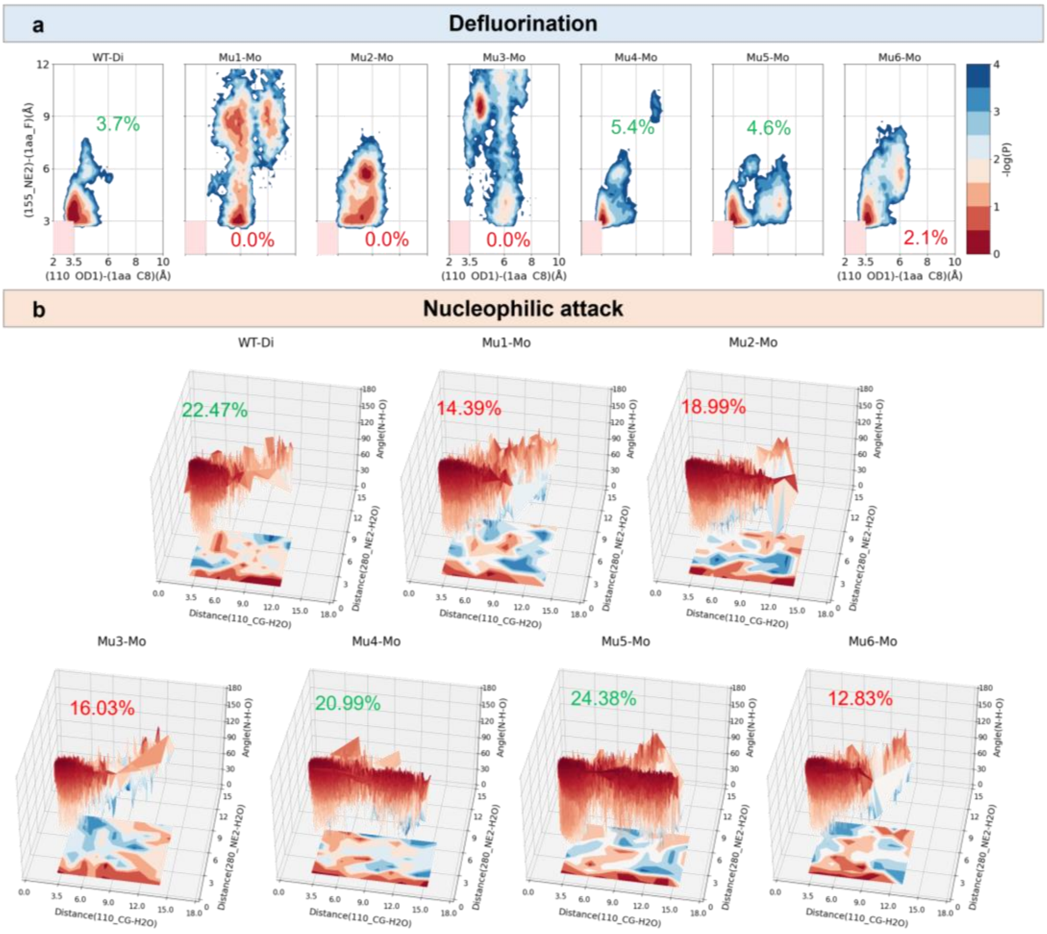

**Figure S6. Ratios of active conformations of WT and mutants during defluorination and nucleophilic attack. a,** Conformational distributions of **WT** and mutants in the defluorination process. **b,** Conformational distributions of **WT** and mutants during the nucleophilic attack.

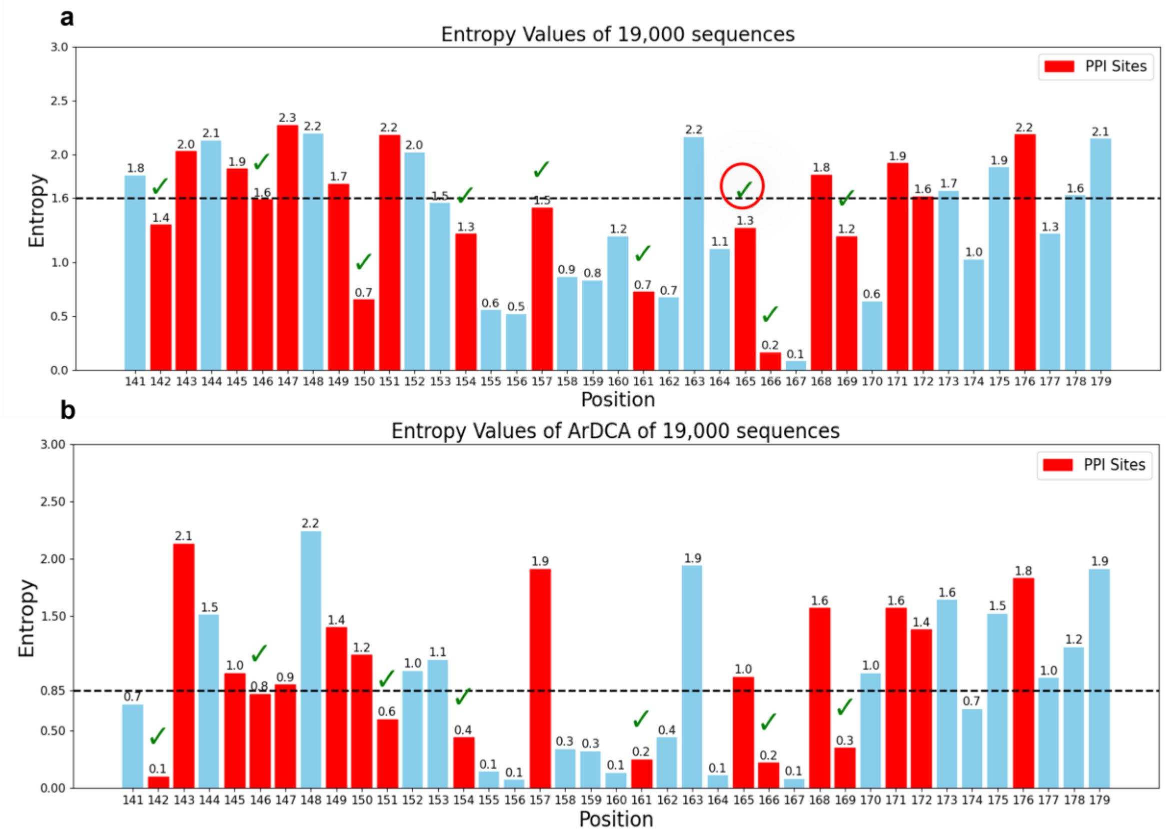

**Figure S7. Entropies of interface sites obtained by different methods. a,** PSSM is obtained by directly analyzed 19,000 homologous sequences and then the entropy of each site is calculated, where L165 was considered to be an active site and cannot be mutated. **b,** 19,000 homologous sequences are input into ArDCA and mutation prediction is performed to obtain PSSM, and the entropy of each site is calculated. It can be seen that the entropy obtained directly using the 19,000 sequences has a low dispersion and it is difficult to distinguish active sites, while the entropy obtained with the help of ArDCA can quickly identify potential active sites.

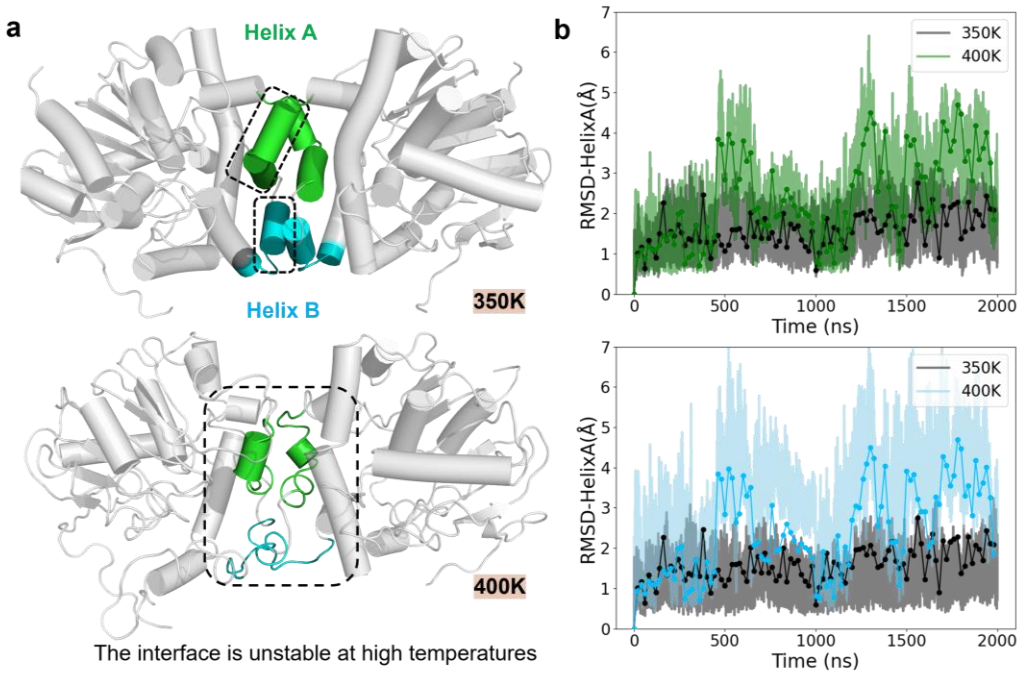

**Figure S8. RMSD of the dimer interface at different temperatures. a,** The main conformations of the FAcD interface at different temperatures, with Helix A and Helix B labeled. **b,** RMSD of the Helix A and Helix B in the FAcD interface at different temperatures.

**Table S1. ProteinMPNN scores of the monomer predicted by AF2.** All scores were subtracted from the wild type, which had a score of 0. Amino acid scores > 0 for stabilizing favorable mutations and < 0 for unfavorable mutations. The color gradient indicates the MPNN score, with greener colors representing higher scores and suggesting more favorable mutations. Conversely, redder colors indicate lower scores and suggest less favorable mutations.

| Index | Wt AA | A | C | D | E | F | G | H | I | K | L | M | N | P | Q | R | S | T | V | W | Y | MAX Score | MAX AA |
|---|---|---|---|---|---|---|---|---|---|---|---|---|---|---|---|---|---|---|---|---|---|---|---|
| **141** | Y | -1.32 | -2.79 | -0.50 | -0.06 | -0.53 | -1.89 | -0.87 | -0.65 | -2.24 | -1.20 | -1.33 | -1.28 | -3.11 | -1.02 | -1.63 | -1.55 | -1.17 | -0.37 | -1.50 | 0.00 | 0 | Y |
| **142** | W | -0.18 | -2.23 | -2.00 | -0.86 | -1.16 | -2.26 | 0.62 | -0.44 | 0.71 | -0.52 | -0.54 | -1.48 | -2.91 | -0.09 | 0.70 | -1.29 | -1.65 | -0.06 | 0.00 | -1.06 | 0.71 | K |
| **143** | Q | 0.71 | -2.36 | -1.11 | -0.31 | -3.02 | -2.53 | -2.15 | -1.78 | 0.65 | -1.04 | -1.32 | -0.68 | -3.31 | 0.00 | 0.19 | 0.02 | -0.80 | -1.48 | -3.04 | -2.51 | 0.71 | A |
| **144** | R | -1.57 | -4.19 | -3.64 | -3.43 | -3.54 | -3.03 | -1.45 | -3.47 | -0.69 | -0.50 | -2.49 | -1.14 | -4.23 | -2.01 | 0.00 | -1.70 | -2.12 | -3.74 | -4.19 | -3.46 | 0 | R |
| **145** | M | -0.19 | -1.03 | -1.51 | -1.45 | 0.53 | -1.33 | -0.84 | -1.42 | -0.33 | -0.23 | 0.00 | 0.14 | -2.67 | -0.17 | 0.55 | 0.04 | -0.95 | -1.35 | 1.13 | 0.41 | 1.13 | W |
| **146** | N | -4.42 | -4.31 | 0.28 | -3.97 | -4.20 | -3.00 | -3.26 | -4.31 | -4.09 | -4.37 | -4.38 | 0.00 | -4.63 | -4.18 | -3.88 | -3.14 | -2.33 | -4.24 | -4.52 | -4.16 | 0.28 | D |
| **147** | R | 1.69 | -1.27 | 0.49 | 0.99 | -0.41 | 1.07 | -0.75 | -0.59 | 0.54 | -0.23 | -0.93 | -0.05 | 1.56 | 0.12 | 0.00 | 1.57 | 0.70 | 0.11 | -0.98 | -0.68 | 1.69 | A |
| **148** | A | 0.00 | -2.90 | 0.00 | 0.26 | -1.21 | -0.64 | -1.35 | -1.30 | -0.23 | -0.99 | -1.94 | -0.83 | -1.96 | -0.54 | -0.26 | -0.12 | 0.06 | -0.32 | -2.10 | -1.18 | 0.26 | E |
| **149** | Y | 0.19 | -1.09 | -0.81 | -0.29 | 0.24 | -1.01 | -0.77 | 1.24 | -0.72 | 1.68 | 2.31 | -0.09 | -0.88 | 0.20 | 0.07 | 0.17 | 1.37 | 2.30 | 1.59 | 0.00 | 2.31 | M |
| **150** | A | 0.00 | -2.01 | -1.28 | -0.29 | -1.47 | -1.44 | -1.05 | -0.64 | 0.67 | 0.68 | -0.34 | -0.31 | -2.52 | 0.77 | 0.62 | -0.05 | -0.33 | -0.74 | -1.54 | -1.14 | 0.77 | Q |
| **151** | L | 1.49 | -1.99 | 0.35 | 1.51 | -1.17 | 0.21 | -0.56 | -0.92 | 1.39 | 0.00 | -0.85 | 0.25 | -2.20 | 1.16 | 0.86 | 0.85 | 0.37 | -0.47 | -1.15 | -1.20 | 1.51 | E |
| **152** | K | -0.05 | -2.86 | -1.24 | -0.57 | -1.11 | -1.72 | -1.10 | -1.91 | 0.00 | 0.93 | -0.95 | -0.92 | -3.05 | -0.31 | 0.38 | -0.55 | -0.95 | -1.87 | -2.48 | -1.22 | 0.93 | L |
| **153** | I | -1.33 | -1.84 | -1.98 | -1.50 | -1.47 | -2.38 | -1.82 | 0.00 | -2.17 | -0.85 | -1.06 | -1.15 | -3.46 | -0.92 | -1.28 | -1.35 | -0.98 | -0.29 | -2.54 | -1.70 | 0 | I |
| **154** | Y | 0.71 | -2.00 | -1.49 | -1.64 | 0.49 | 0.57 | -0.24 | -2.00 | -1.32 | -1.11 | -1.73 | 0.01 | -2.04 | -0.79 | -0.03 | 0.24 | -1.62 | -1.47 | 0.79 | 0.00 | 0.79 | W |
| **155** | H | 1.05 | 0.39 | 2.35 | 4.25 | 3.17 | 0.37 | 2.41 | 0.98 | 0.05 | 2.40 | 1.06 | 1.01 | 0.65 | 2.06 | 0.18 | 1.09 | 0.94 | 0.75 | 2.73 | 3.17 | 4.25 | E |
| **156** | W | -0.91 | -1.80 | -1.09 | -1.09 | 0.77 | -1.47 | -0.42 | 0.90 | -0.98 | 1.83 | 0.88 | -0.75 | -2.41 | -0.26 | 0.01 | -1.06 | -0.95 | -0.45 | 0.00 | 0.49 | 1.83 | L |
| **157** | S | 0.16 | -1.90 | -1.39 | -1.07 | -0.52 | -0.04 | -1.31 | 0.11 | -0.60 | 1.48 | -0.38 | -0.67 | -1.99 | -0.19 | -0.27 | 0.00 | 0.37 | 0.30 | -1.03 | -1.56 | 1.48 | L |
| **158** | F | -2.13 | -3.70 | -3.95 | -3.74 | 0.00 | -2.89 | -3.43 | -2.04 | -3.87 | -1.03 | -2.62 | -3.53 | -4.59 | -3.24 | -3.14 | -3.07 | -2.67 | -1.38 | -2.18 | -1.96 | 0 | F |
| **159** | L | -4.46 | -4.90 | -4.72 | -4.65 | -2.68 | -4.85 | -3.81 | -4.25 | -4.81 | 0.00 | -2.00 | -4.27 | -5.25 | -4.45 | -4.25 | -4.75 | -5.00 | -5.09 | -3.84 | -2.96 | 0 | L |
| **160** | A | 0.00 | -4.79 | -4.85 | -4.91 | -4.67 | -4.16 | -4.86 | -4.67 | -5.08 | -4.88 | -4.89 | -4.43 | -4.92 | -4.81 | -4.57 | -0.37 | -4.93 | -5.01 | -4.87 | -4.91 | 0 | A |
| **161** | Q | 0.08 | -1.90 | -2.79 | -1.97 | -1.18 | -2.51 | -2.12 | -2.14 | -1.67 | 1.87 | 0.67 | -2.61 | -3.03 | 0.00 | -1.13 | -1.65 | -1.47 | -2.46 | -1.41 | -1.97 | 1.87 | L |
| **162** | P | -2.35 | -5.20 | -2.84 | -2.50 | -4.84 | -4.08 | -4.21 | -4.56 | -3.15 | -4.11 | -4.70 | -4.04 | 0.00 | -3.53 | -3.28 | -3.09 | -3.65 | -4.18 | -4.95 | -4.75 | 0 | P |
| **163** | A | 0.00 | -3.48 | -1.60 | -2.12 | -1.37 | -2.12 | -1.04 | -2.35 | -1.94 | -0.66 | -2.37 | -2.29 | -0.67 | -1.89 | -1.32 | -0.92 | -2.70 | -3.02 | -2.80 | -2.23 | 0 | A |
| **164** | P | -5.02 | -5.41 | -5.00 | -5.29 | -5.24 | -5.05 | -5.19 | -5.45 | -5.36 | -5.26 | -5.44 | -5.13 | 0.00 | -5.39 | -5.39 | -4.95 | -5.15 | -5.39 | -5.40 | -5.48 | 0 | P |
| **165** | L | -0.31 | -2.34 | -2.11 | -1.59 | 2.29 | -1.35 | -0.94 | 0.47 | -1.58 | 0.00 | -1.34 | -2.11 | -2.50 | -1.63 | -1.62 | -1.69 | -0.51 | 1.00 | -0.52 | 1.18 | 2.29 | F |
| **166** | P | 1.19 | -3.22 | -3.70 | -3.61 | -3.56 | -3.57 | -3.90 | -3.78 | -3.73 | -3.58 | -3.31 | -3.59 | 0.00 | -3.63 | -3.21 | -1.83 | -2.92 | -3.48 | -3.38 | -3.49 | 1.19 | A |
| **167** | E | -2.79 | -4.56 | -2.68 | 0.00 | -4.40 | -2.90 | -4.38 | -2.28 | -4.45 | -0.50 | -2.73 | -3.83 | -3.75 | -1.32 | -3.79 | -2.62 | -3.16 | -2.38 | -4.59 | -4.24 | 0 | E |
| **168** | N | 1.28 | -1.84 | 0.24 | 0.16 | -1.07 | -0.69 | -1.05 | 0.11 | 0.71 | 1.01 | -0.57 | 0.00 | -1.89 | 0.28 | 0.32 | 0.50 | 1.07 | 0.63 | -1.62 | -1.53 | 1.28 | A |
| **169** | L | -0.14 | -2.34 | -2.63 | -2.21 | 0.75 | -2.18 | -1.79 | 1.13 | -1.95 | 0.00 | -1.11 | -2.37 | -2.64 | -1.89 | -1.81 | -1.83 | -0.91 | 1.19 | -0.65 | -0.14 | 1.19 | V |
| **170** | L | -5.12 | -5.25 | -5.36 | -5.12 | -3.72 | -5.35 | -5.34 | -2.13 | -5.24 | 0.00 | -2.99 | -5.21 | -5.37 | -4.87 | -5.08 | -5.18 | -5.08 | -4.10 | -3.61 | -3.24 | 0 | L |
| **171** | G | 1.53 | -2.21 | -1.05 | -1.36 | -2.41 | 0.00 | -2.12 | -2.30 | -1.20 | 0.30 | -0.49 | 0.38 | -2.56 | -0.89 | -0.97 | 0.76 | -2.07 | -2.62 | -2.73 | -2.54 | 1.53 | A |
| **172** | G | 1.46 | -0.58 | -0.44 | 0.50 | 1.52 | 0.00 | 0.09 | 2.76 | 1.34 | 2.99 | 1.01 | -0.12 | -0.72 | 1.39 | 1.19 | 0.19 | 0.63 | 2.58 | 0.23 | 0.33 | 2.99 | L |
| **173** | D | -2.65 | -4.66 | 0.00 | -3.64 | -4.54 | -4.26 | -3.53 | -4.89 | -4.23 | -4.62 | -4.51 | -0.51 | -4.86 | -3.84 | -3.99 | -3.07 | -4.27 | -4.59 | -4.87 | -4.69 | 0 | D |
| **174** | P | -2.77 | -5.36 | -5.44 | -5.34 | -5.16 | -5.39 | -5.30 | -5.59 | -5.53 | -5.25 | -5.42 | -5.33 | 0.00 | -5.33 | -5.25 | -3.66 | -5.27 | -5.38 | -5.04 | -5.05 | 0 | P |
| **175** | D | -2.62 | -5.36 | 0.00 | -1.22 | -4.92 | -2.97 | -4.44 | -4.56 | -5.14 | -3.53 | -4.80 | -4.35 | -5.10 | -4.15 | -4.34 | -4.30 | -4.00 | -3.73 | -4.77 | -4.56 | 0 | D |
| **176** | F | 3.03 | -1.85 | -0.76 | -0.46 | 0.00 | 1.04 | 0.46 | -1.35 | 0.88 | -0.19 | -1.53 | -0.83 | -1.70 | -0.20 | 1.19 | 1.04 | 0.20 | -0.56 | -0.95 | 0.11 | 3.03 | A |
| **177** | Y | -0.74 | -2.74 | -2.81 | -2.65 | 1.19 | -3.07 | -1.88 | -0.99 | -3.10 | -1.33 | -1.52 | -3.01 | -3.42 | -2.51 | -3.00 | -2.41 | -2.22 | -0.40 | -0.04 | 0.00 | 1.19 | F |
| **178** | V | -4.36 | -2.68 | -4.88 | -4.50 | -5.13 | -4.96 | -5.19 | -1.88 | -5.25 | -3.47 | -4.18 | -3.52 | -5.15 | -3.97 | -4.87 | -4.59 | -3.77 | 0.00 | -5.10 | -5.12 | 0 | V |
| **179** | K | 1.54 | -0.43 | 3.53 | 2.30 | -0.80 | 0.64 | -0.24 | -0.66 | 0.00 | 1.18 | -0.54 | 2.14 | -0.73 | 0.48 | 0.85 | 2.10 | 2.25 | -0.02 | -0.86 | -0.55 | 3.53 | D |

**Table S2. ProteinMPNN scores of the dimer predicted by AF2.** All scores were subtracted from the wild type, which had a score of 0. Amino acid scores > 0 for stabilizing favorable mutations and < 0 for unfavorable mutations. The color gradient indicates the MPNN score, with greener colors representing higher scores and suggesting more favorable mutations. Conversely, redder colors indicate lower scores and suggest less favorable mutations.

| Index | Wt AA | A | C | D | E | F | G | H | I | K | L | M | N | P | Q | R | S | T | V | W | Y | MAX Score | MAX AA |
|---|---|---|---|---|---|---|---|---|---|---|---|---|---|---|---|---|---|---|---|---|---|---|---|
| **141** | Y | -1.49 | -2.61 | -0.48 | 0.48 | -1.03 | -2.18 | -1.81 | 0.81 | -2.39 | -0.71 | -0.37 | -1.46 | -2.90 | -0.58 | -1.66 | -1.76 | -0.65 | 0.97 | -0.28 | 0.00 | 0.97 | V |
| **142** | W | -3.52 | -4.74 | -3.98 | -3.78 | -2.32 | -4.44 | -2.18 | -4.82 | -4.35 | -1.39 | -3.42 | -3.48 | -5.00 | -2.66 | -4.29 | -4.15 | -4.80 | -4.86 | 0.00 | -1.99 | 0 | W |
| **143** | Q | -5.01 | -5.59 | -5.37 | -5.44 | -5.68 | -5.61 | -5.69 | -5.64 | -5.54 | -5.63 | -5.56 | -5.18 | -5.74 | 0.00 | -5.54 | -5.24 | -5.46 | -5.62 | -5.65 | -5.55 | 0 | Q |
| **144** | R | -5.75 | -5.66 | -5.76 | -5.77 | -5.68 | -5.60 | -5.72 | -5.68 | -5.70 | -5.73 | -5.73 | -5.62 | -5.75 | -5.74 | 0.00 | -5.68 | -5.65 | -5.68 | -5.63 | -5.70 | 0 | R |
| **145** | M | -5.82 | -5.75 | -5.84 | -5.82 | -5.69 | -5.84 | -5.76 | -5.73 | -5.73 | -6.07 | 0.00 | -5.70 | -5.78 | -5.83 | -5.73 | -5.76 | -5.81 | -5.81 | -5.70 | -5.48 | 0 | M |
| **146** | N | -5.61 | -5.51 | -5.19 | -5.64 | -5.66 | -5.60 | -5.62 | -5.54 | -5.70 | -5.70 | -5.62 | 0.00 | -5.63 | -5.59 | -5.65 | -5.61 | -5.64 | -5.58 | -5.56 | -5.62 | 0 | N |
| **147** | R | -5.78 | -5.75 | -5.82 | -5.81 | -5.80 | -5.78 | -5.48 | -5.74 | -5.61 | -5.70 | -5.77 | -5.71 | -5.81 | -5.80 | 0.00 | -5.84 | -5.82 | -5.81 | -5.64 | -5.75 | 0 | R |
| **148** | A | 0.00 | -2.89 | 1.52 | 1.49 | -1.79 | -1.13 | -1.21 | -2.46 | -0.62 | -2.13 | -2.64 | -0.63 | -2.58 | -0.46 | -1.11 | -0.64 | -0.87 | -1.69 | -2.18 | -1.31 | 1.52 | D |
| **149** | Y | -5.60 | -5.62 | -5.72 | -5.47 | -5.81 | -5.69 | -5.75 | -5.61 | -5.68 | -5.51 | -5.50 | -5.59 | -5.69 | -5.64 | -5.59 | -5.56 | -5.63 | -5.63 | -5.31 | 0.00 | 0 | Y |
| **150** | A | 0.00 | -5.53 | -5.62 | -5.72 | -5.55 | -5.61 | -5.57 | -5.58 | -5.68 | -5.63 | -5.56 | -5.53 | -5.72 | -5.62 | -5.57 | -5.56 | -5.54 | -5.59 | -5.55 | -5.57 | 0 | A |
| **151** | L | -5.58 | -5.62 | -5.54 | -5.52 | -5.61 | -5.52 | -5.51 | -5.66 | -5.63 | 0.00 | -5.61 | -5.55 | -5.62 | -5.48 | -5.61 | -5.52 | -5.57 | -5.67 | -5.61 | -5.52 | 0 | L |
| **152** | K | 0.07 | -3.49 | -1.42 | -0.75 | -3.46 | -2.02 | -2.61 | -3.43 | 0.00 | -1.13 | -2.16 | -1.11 | -3.55 | -0.63 | -0.11 | -0.43 | -1.12 | -3.33 | -3.62 | -3.09 | 0.07 | A |
| **153** | I | 0.24 | -0.65 | -1.51 | -0.77 | -2.29 | -1.80 | -1.93 | 0.00 | -1.99 | -0.33 | -0.41 | -0.80 | -3.00 | -0.72 | -0.54 | -0.15 | -0.60 | 0.08 | -2.00 | -1.72 | 0.24 | A |
| **154** | Y | -5.57 | -5.56 | -5.58 | -5.49 | -5.51 | -5.63 | -5.55 | -5.65 | -5.43 | -5.67 | -5.56 | -5.51 | -5.59 | -5.55 | -5.68 | -5.48 | -5.54 | -5.66 | -5.47 | 0.00 | 0 | Y |
| **155** | H | -5.71 | -5.63 | -5.60 | -5.65 | -5.70 | -5.77 | 0.00 | -5.68 | -5.80 | -5.77 | -5.75 | -5.57 | -5.74 | -5.25 | -5.65 | -5.71 | -5.70 | -5.72 | -5.62 | -5.71 | 0 | H |
| **156** | W | -1.09 | -2.19 | -0.23 | -0.73 | 0.66 | -1.65 | -0.69 | -1.10 | -1.77 | 1.15 | -0.06 | -0.50 | -2.54 | -0.47 | -0.88 | -1.24 | -1.47 | -1.79 | 0.00 | 0.78 | 1.15 | L |
| **157** | S | -5.59 | -5.63 | -5.90 | -5.80 | -5.59 | -5.81 | -5.54 | -5.61 | -5.71 | -5.69 | -5.68 | -5.63 | -5.76 | -5.73 | -5.75 | 0.00 | -5.28 | -5.79 | -5.71 | -5.68 | 0 | S |
| **158** | F | -5.72 | -5.63 | -5.67 | -5.70 | 0.00 | -5.67 | -5.75 | -5.63 | -5.65 | -5.37 | -5.71 | -5.62 | -5.67 | -5.65 | -5.68 | -5.75 | -5.60 | -5.67 | -5.62 | -5.76 | 0 | F |
| **159** | L | -5.33 | -5.28 | -5.10 | -4.88 | -2.91 | -5.44 | -4.22 | -5.00 | -5.07 | 0.00 | -3.57 | -4.64 | -5.46 | -4.94 | -5.02 | -5.30 | -5.43 | -5.64 | -4.34 | -3.64 | 0 | L |
| **160** | A | 0.00 | -4.70 | -4.87 | -4.76 | -4.57 | -4.89 | -4.96 | -4.40 | -4.86 | -4.88 | -4.63 | -4.75 | -4.79 | -4.89 | -4.78 | -0.38 | -4.75 | -4.82 | -4.83 | -4.70 | 0 | A |
| **161** | Q | -3.62 | -5.88 | -6.42 | -5.59 | -6.11 | -5.76 | -5.72 | -6.13 | -4.24 | -3.10 | -3.77 | -5.87 | -6.12 | 0.00 | -4.21 | -5.39 | -5.38 | -5.99 | -5.41 | -6.06 | 0 | Q |
| **162** | P | -2.59 | -5.26 | -3.03 | -2.48 | -5.21 | -4.67 | -4.26 | -5.23 | -2.90 | -4.93 | -5.18 | -4.57 | 0.00 | -3.92 | -3.32 | -3.71 | -4.24 | -4.84 | -5.09 | -4.86 | 0 | P |
| **163** | A | 0.00 | -3.60 | -2.20 | -2.59 | -1.69 | -2.84 | -0.02 | -2.43 | -1.42 | -1.45 | -3.05 | -2.46 | -1.67 | -1.58 | -0.71 | -1.30 | -2.44 | -3.02 | -1.92 | -1.56 | 0 | A |
| **164** | P | -5.42 | -5.45 | -5.48 | -5.45 | -5.42 | -5.46 | -5.42 | -5.42 | -5.48 | -5.46 | -5.44 | -5.46 | 0.00 | -5.51 | -5.51 | -5.48 | -5.42 | -5.49 | -5.45 | -5.45 | 0 | P |
| **165** | L | -3.42 | -3.72 | -3.65 | -3.20 | -0.84 | -3.76 | -3.44 | 0.96 | -3.43 | 0.00 | -1.25 | -3.64 | -3.50 | -3.24 | -3.33 | -3.65 | -2.12 | -0.16 | -1.96 | -1.90 | 0.96 | I |
| **166** | P | -5.30 | -5.53 | -5.52 | -5.73 | -5.58 | -5.79 | -5.62 | -5.54 | -5.59 | -5.63 | -5.54 | -5.52 | 0.00 | -5.75 | -5.50 | -5.75 | -5.56 | -5.58 | -5.70 | -5.54 | 0 | P |
| **167** | E | -4.98 | -5.54 | -4.29 | 0.00 | -5.78 | -4.57 | -5.82 | -4.41 | -5.67 | -3.46 | -5.32 | -4.61 | -5.09 | -2.65 | -5.34 | -5.25 | -4.86 | -3.24 | -5.47 | -5.13 | 0 | E |
| **168** | N | -5.51 | -5.72 | -5.70 | -5.69 | -5.70 | -5.68 | -5.73 | -5.68 | -5.35 | -5.77 | -5.77 | 0.00 | -5.70 | -5.50 | -5.43 | -5.73 | -5.57 | -5.61 | -5.66 | -5.69 | 0 | N |
| **169** | L | -5.56 | -5.62 | -5.58 | -5.57 | -5.65 | -5.57 | -5.57 | -5.71 | -5.60 | 0.00 | -5.68 | -5.63 | -5.61 | -5.68 | -5.65 | -5.55 | -5.60 | -5.66 | -5.60 | -5.54 | 0 | L |
| **170** | L | -5.65 | -5.59 | -5.57 | -5.60 | -5.59 | -5.61 | -5.58 | -5.35 | -5.64 | 0.00 | -5.63 | -5.62 | -5.68 | -5.71 | -5.70 | -5.57 | -5.64 | -5.51 | -5.67 | -5.48 | 0 | L |
| **171** | G | -5.52 | -5.60 | -5.64 | -5.72 | -5.53 | 0.00 | -5.55 | -5.58 | -5.62 | -5.47 | -5.48 | -5.49 | -5.79 | -5.53 | -5.65 | -5.60 | -5.64 | -5.60 | -5.56 | -5.53 | 0 | G |
| **172** | G | -5.69 | -5.68 | -5.82 | -5.81 | -5.57 | 0.00 | -5.67 | -5.67 | -5.66 | -5.68 | -5.66 | -5.25 | -5.85 | -5.79 | -5.73 | -5.83 | -5.60 | -5.53 | -5.64 | -5.68 | 0 | G |
| **173** | D | -5.53 | -5.50 | 0.00 | -5.78 | -5.53 | -5.65 | -5.54 | -5.53 | -5.62 | -5.61 | -5.51 | -5.35 | -5.60 | -5.63 | -5.67 | -5.65 | -5.44 | -5.58 | -5.62 | -5.52 | 0 | D |
| **174** | P | -1.89 | -5.16 | -5.18 | -5.11 | -5.05 | -5.56 | -5.17 | -5.34 | -5.48 | -4.90 | -5.27 | -5.16 | 0.00 | -5.26 | -4.15 | -3.08 | -4.26 | -4.22 | -4.72 | -4.87 | 0 | P |
| **175** | D | -3.42 | -5.37 | 0.00 | -1.07 | -5.22 | -4.06 | -5.20 | -4.97 | -5.42 | -4.71 | -5.39 | -4.93 | -5.34 | -5.00 | -4.83 | -4.84 | -4.52 | -4.08 | -5.08 | -4.82 | 0 | D |
| **176** | F | -3.79 | -5.90 | -5.51 | -5.39 | 0.00 | -3.00 | -5.37 | -5.61 | -5.69 | -4.59 | -5.42 | -5.41 | -5.87 | -5.32 | -5.11 | -4.95 | -5.39 | -5.45 | -5.01 | -2.86 | 0 | F |
| **177** | Y | -1.24 | -3.00 | -2.78 | -2.75 | 2.17 | -3.00 | -2.88 | -2.87 | -2.57 | -2.42 | -2.50 | -2.87 | -2.76 | -2.90 | -2.71 | -2.66 | -2.21 | -1.24 | 0.26 | 0.00 | 2.17 | F |
| **178** | V | -4.75 | -3.36 | -4.98 | -4.71 | -4.89 | -5.15 | -4.86 | -0.63 | -5.06 | -2.34 | -4.15 | -4.56 | -5.07 | -4.69 | -4.89 | -4.94 | -4.30 | 0.00 | -4.92 | -4.75 | 0 | V |
| **179** | K | 0.93 | -0.45 | 3.32 | 2.81 | -1.07 | 0.51 | 0.13 | -0.66 | 0.00 | 1.24 | -0.71 | 2.08 | -0.86 | 0.51 | 1.24 | 1.18 | 2.40 | -0.19 | -0.81 | -0.47 | 3.32 | D |

**Table S3. ProteinMPNN scores of the dimeric crystal structure (3R3U).** All scores were subtracted from the wild type, which had a score of 0. Amino acid scores > 0 for stabilizing favorable mutations and < 0 for unfavorable mutations. The color gradient indicates the MPNN score, with greener colors representing higher scores and suggesting more favorable mutations. Conversely, redder colors indicate lower scores and suggest less favorable mutations.

| Index | Wt AA | A | C | D | E | F | G | H | I | K | L | M | N | P | Q | R | S | T | V | W | Y | MAX Score | MAX AA |
|---|---|---|---|---|---|---|---|---|---|---|---|---|---|---|---|---|---|---|---|---|---|---|---|
| 141 | Y | -3.18 | -4.23 | -2.90 | -1.90 | -0.49 | -3.77 | -3.02 | -1.34 | -4.13 | -1.67 | -2.10 | -3.30 | -4.44 | -2.82 | -3.45 | -3.68 | -2.63 | -1.24 | -1.78 | 0.00 | 0 | Y |
| 142 | W | -2.66 | -4.25 | -3.73 | -3.21 | -3.58 | -4.60 | -0.30 | -4.64 | -4.10 | -3.17 | -2.43 | -4.18 | -4.79 | -2.02 | -4.16 | -3.95 | -4.84 | -4.64 | 0.00 | -3.53 | 0 | W |
| 143 | Q | -5.50 | -5.60 | -5.62 | -5.68 | -5.56 | -5.66 | -5.74 | -5.62 | -5.57 | -5.49 | -5.55 | -5.51 | -5.72 | 0.00 | -5.65 | -5.58 | -5.56 | -5.62 | -5.57 | -5.57 | 0 | Q |
| 144 | R | -5.81 | -5.94 | -5.93 | -6.09 | -6.09 | -5.84 | -5.41 | -6.06 | -5.43 | -5.44 | -6.12 | -5.60 | -6.07 | -5.87 | 0.00 | -6.10 | -5.80 | -5.89 | -6.07 | -6.05 | 0 | R |
| 145 | M | -5.85 | -5.85 | -5.87 | -5.86 | -5.41 | -5.86 | -5.75 | -5.78 | -5.84 | -5.86 | 0.00 | -5.83 | -5.86 | -5.88 | -5.82 | -5.78 | -5.90 | -5.97 | -5.68 | -5.68 | 0 | M |
| 146 | N | -5.70 | -5.65 | -4.86 | -5.74 | -5.75 | -5.65 | -5.60 | -5.62 | -5.83 | -5.78 | -5.72 | 0.00 | -5.71 | -5.69 | -5.80 | -5.79 | -5.64 | -5.62 | -5.65 | -5.69 | 0 | N |
| 147 | R | -5.80 | -5.75 | -5.84 | -5.83 | -5.79 | -5.79 | -5.46 | -5.75 | -5.61 | -5.70 | -5.78 | -5.77 | -5.79 | -5.82 | 0.00 | -5.84 | -5.80 | -5.80 | -5.62 | -5.76 | 0 | R |
| 148 | A | 0.00 | -2.78 | 1.44 | 1.67 | -2.52 | -1.26 | -1.62 | -2.34 | -0.25 | -2.10 | -2.62 | -0.73 | -2.29 | -0.07 | -0.85 | -0.63 | -0.56 | -1.48 | -2.29 | -2.03 | 1.67 | E |
| 149 | Y | -5.68 | -5.61 | -5.63 | -5.48 | -5.53 | -5.61 | -5.58 | -5.40 | -5.79 | -5.33 | -5.23 | -5.48 | -5.67 | -5.51 | -5.60 | -5.60 | -5.55 | -5.64 | -5.23 | 0.00 | 0 | Y |
| 150 | A | 0.00 | -5.54 | -5.64 | -5.76 | -5.57 | -5.59 | -5.57 | -5.64 | -5.74 | -5.66 | -5.62 | -5.51 | -5.73 | -5.61 | -5.61 | -5.45 | -5.51 | -5.58 | -5.59 | -5.64 | 0 | A |
| 151 | L | -5.60 | -5.64 | -5.59 | -5.58 | -5.66 | -5.56 | -5.55 | -5.67 | -5.65 | 0.00 | -5.70 | -5.59 | -5.64 | -5.45 | -5.63 | -5.53 | -5.57 | -5.72 | -5.63 | -5.57 | 0 | L |
| 152 | K | -1.38 | -4.36 | -2.47 | -1.65 | -3.41 | -3.53 | -2.62 | -4.12 | 0.00 | -1.24 | -2.79 | -2.65 | -4.26 | -1.27 | -0.17 | -1.82 | -2.65 | -4.12 | -4.08 | -3.07 | 0 | K |
| 153 | I | -1.37 | -1.73 | -1.55 | -0.98 | -1.00 | -2.18 | -2.35 | 0.00 | -3.27 | -0.32 | -1.19 | -1.23 | -3.73 | -1.46 | -2.48 | -1.95 | -1.61 | -0.50 | -1.45 | -1.12 | 0 | I |
| 154 | Y | -5.57 | -5.55 | -5.60 | -5.48 | -5.51 | -5.62 | -5.57 | -5.63 | -5.43 | -5.66 | -5.56 | -5.52 | -5.58 | -5.57 | -5.66 | -5.50 | -5.54 | -5.65 | -5.50 | 0.00 | 0 | Y |
| 155 | H | -5.72 | -5.66 | -5.53 | -5.63 | -5.66 | -5.79 | 0.00 | -5.68 | -5.79 | -5.76 | -5.78 | -5.60 | -5.75 | -5.44 | -5.66 | -5.74 | -5.71 | -5.74 | -5.68 | -5.71 | 0 | H |
| 156 | W | -1.71 | -2.61 | -1.81 | -1.82 | 0.01 | -2.54 | -1.34 | -1.44 | -1.93 | 1.33 | 0.32 | -1.37 | -2.91 | -1.15 | -1.00 | -2.05 | -1.83 | -2.13 | 0.00 | 0.05 | 1.33 | L |
| 157 | S | -5.55 | -5.57 | -5.90 | -5.81 | -5.58 | -5.80 | -5.63 | -5.62 | -5.71 | -5.69 | -5.69 | -5.64 | -5.78 | -5.70 | -5.72 | 0.00 | -5.22 | -5.79 | -5.70 | -5.68 | 0 | S |
| 158 | F | -5.72 | -5.62 | -5.66 | -5.70 | 0.00 | -5.67 | -5.73 | -5.61 | -5.63 | -5.39 | -5.68 | -5.62 | -5.66 | -5.65 | -5.67 | -5.73 | -5.60 | -5.66 | -5.56 | -5.72 | 0 | F |
| 159 | L | -5.08 | -5.26 | -4.78 | -4.41 | -2.27 | -5.35 | -3.82 | -4.77 | -5.02 | 0.00 | -3.27 | -4.64 | -5.22 | -4.81 | -4.75 | -5.08 | -5.25 | -5.46 | -3.57 | -2.66 | 0 | L |
| 160 | A | 0.00 | -5.12 | -5.17 | -5.21 | -4.93 | -5.07 | -5.27 | -4.71 | -5.24 | -5.17 | -5.07 | -5.18 | -5.07 | -5.28 | -5.09 | -1.40 | -5.08 | -5.16 | -5.13 | -4.97 | 0 | A |
| 161 | Q | -2.58 | -5.18 | -6.10 | -5.30 | -5.82 | -5.59 | -5.47 | -5.87 | -3.72 | -3.49 | -3.68 | -5.64 | -5.86 | 0.00 | -3.18 | -4.75 | -4.62 | -5.34 | -5.21 | -5.69 | 0 | Q |
| 162 | P | -2.34 | -5.27 | -2.70 | -2.04 | -5.16 | -4.70 | -4.42 | -5.23 | -3.22 | -4.90 | -5.18 | -4.62 | 0.00 | -3.81 | -3.67 | -3.82 | -4.22 | -4.82 | -5.15 | -4.91 | 0 | P |
| 163 | A | 0.00 | -3.50 | -1.71 | -2.34 | -1.69 | -2.50 | -0.60 | -1.84 | -1.64 | -0.38 | -2.38 | -2.22 | -1.88 | -1.56 | -0.93 | -1.35 | -2.60 | -3.33 | -2.88 | -2.07 | 0 | A |
| 164 | P | -5.41 | -5.46 | -5.48 | -5.47 | -5.42 | -5.44 | -5.41 | -5.43 | -5.50 | -5.46 | -5.42 | -5.47 | 0.00 | -5.51 | -5.50 | -5.49 | -5.42 | -5.49 | -5.45 | -5.45 | 0 | P |
| 165 | L | -2.76 | -3.33 | -3.15 | -2.85 | 1.02 | -3.20 | -2.42 | 0.84 | -3.14 | 0.00 | -0.49 | -3.06 | -3.08 | -2.69 | -2.82 | -3.11 | -1.64 | 0.11 | -0.72 | -0.43 | 1.02 | F |
| 166 | P | -5.08 | -5.46 | -5.46 | -5.67 | -5.57 | -5.76 | -5.58 | -5.47 | -5.54 | -5.58 | -5.51 | -5.45 | 0.00 | -5.73 | -5.46 | -5.68 | -5.53 | -5.62 | -5.67 | -5.55 | 0 | P |
| 167 | E | -3.03 | -4.81 | -3.07 | 0.00 | -4.69 | -2.97 | -4.51 | -2.75 | -5.00 | -0.66 | -3.34 | -4.37 | -4.53 | -1.77 | -4.30 | -3.50 | -4.07 | -3.21 | -4.40 | -4.10 | 0 | E |
| 168 | N | -5.38 | -5.65 | -5.62 | -5.63 | -5.66 | -5.64 | -5.66 | -5.64 | -5.44 | -5.75 | -5.73 | 0.00 | -5.67 | -5.46 | -5.42 | -5.64 | -5.60 | -5.61 | -5.64 | -5.64 | 0 | N |
| 169 | L | -5.57 | -5.57 | -5.56 | -5.57 | -5.60 | -5.54 | -5.56 | -5.77 | -5.59 | 0.00 | -5.58 | -5.62 | -5.58 | -5.67 | -5.63 | -5.53 | -5.57 | -5.65 | -5.59 | -5.50 | 0 | L |
| 170 | L | -5.67 | -5.58 | -5.55 | -5.58 | -5.54 | -5.58 | -5.58 | -5.43 | -5.61 | 0.00 | -5.62 | -5.59 | -5.63 | -5.70 | -5.67 | -5.55 | -5.64 | -5.65 | -5.61 | -5.41 | 0 | L |
| 171 | G | -5.43 | -5.65 | -5.73 | -5.78 | -5.57 | 0.00 | -5.63 | -5.65 | -5.49 | -5.51 | -5.60 | -5.50 | -5.88 | -5.55 | -5.60 | -5.73 | -5.69 | -5.67 | -5.60 | -5.63 | 0 | G |
| 172 | G | -5.59 | -5.61 | -5.70 | -5.63 | -5.54 | 0.00 | -5.59 | -5.61 | -5.72 | -5.63 | -5.62 | -5.54 | -5.77 | -5.70 | -5.67 | -5.68 | -5.57 | -5.57 | -5.59 | -5.63 | 0 | G |
| 173 | D | -5.58 | -5.60 | 0.00 | -5.72 | -5.58 | -5.73 | -5.60 | -5.61 | -5.60 | -5.66 | -5.57 | -5.29 | -5.69 | -5.59 | -5.67 | -5.68 | -5.56 | -5.65 | -5.70 | -5.61 | 0 | D |
| 174 | P | -2.85 | -5.25 | -5.51 | -5.47 | -5.41 | -5.58 | -5.24 | -5.65 | -5.76 | -5.57 | -5.69 | -5.38 | 0.00 | -5.41 | -5.53 | -4.19 | -5.35 | -5.28 | -5.07 | -5.33 | 0 | P |
| 175 | D | -2.73 | -5.28 | 0.00 | -0.80 | -4.97 | -3.26 | -4.68 | -4.61 | -5.30 | -3.37 | -4.85 | -4.78 | -5.07 | -4.11 | -4.44 | -4.50 | -4.09 | -3.90 | -4.88 | -4.78 | 0 | D |
| 176 | F | -1.97 | -5.79 | -5.30 | -5.19 | 0.00 | -3.79 | -5.04 | -5.66 | -5.64 | -4.28 | -5.15 | -5.51 | -5.70 | -5.07 | -4.43 | -4.27 | -5.00 | -5.14 | -4.95 | -3.11 | 0 | F |
| 177 | Y | -2.23 | -3.33 | -2.98 | -3.02 | 1.00 | -3.51 | -3.04 | -2.12 | -2.92 | -2.60 | -3.05 | -3.23 | -3.21 | -3.31 | -3.09 | -3.11 | -2.69 | 0.42 | -0.16 | 0.00 | 1 | F |
| 178 | V | -4.60 | -2.86 | -5.09 | -4.61 | -5.35 | -5.16 | -5.50 | -2.51 | -5.48 | -4.31 | -4.55 | -3.41 | -5.41 | -4.22 | -5.10 | -4.98 | -3.61 | 0.00 | -4.91 | -5.21 | 0 | V |
| 179 | K | 1.80 | 0.26 | 5.40 | 3.81 | -0.03 | 1.12 | 0.48 | 0.07 | 0.00 | 1.51 | 0.20 | 2.16 | 0.14 | 0.82 | 0.70 | 1.61 | 1.78 | 0.41 | 0.17 | 0.17 | 5.4 | D |

**Table S4. Contact Frequency of Interface Amino Acids (Y141-K179).** The total frequency of all interactions at the interface, including hydrogen bonds, salt bridges, pi-pi interactions, van der Waals, and hydrophobic interactions, was analyzed over 3 × 1 μs molecular dynamics simulations using GetContacts. Amino acids with a contact frequency greater than 0.6 are highlighted, among which Y149 and L165 were not identified.

| ChainA | ChainB | ContactFrequency | Total | $MAX\ (m_i)$ |
|---|---|---|---|---|
| L169 | L169 | 0.988 | W142 | 0.71 |
| Q161 | L151 | 0.982 | M145 | 1.13 |
| L151 | Q161 | 0.982 | N146 | 0.28 |
| Q161 | Y154 | 0.978 | R147 | 1.69 |
| Y154 | Q161 | 0.977 | A150 | 0.77 |
| F158 | F158 | 0.919 | L151 | 1.51 |
| S157 | A150 | 0.914 | Y154 | 0.79 |
| A150 | S157 | 0.891 | S157 | 1.48 |
| W142 | R147 | 0.889 | F158 | 0 |
| R147 | W142 | 0.852 | A160 | 0 |
| M145 | M145 | 0.806 | Q161 | 1.87 |
| N168 | F176 | 0.722 | N168 | 1.28 |
| M145 | N146 | 0.697 | L169 | 1.19 |
| F176 | N168 | 0.691 | F176 | 3.03 |
| N146 | M145 | 0.636 | | |
| A160 | L151 | 0.635 | | |
| L151 | A160 | 0.613 | | |
